# The structure of the SufS-SufE complex reveals interactions driving protected persulfide transfer in iron-sulfur cluster biogenesis

**DOI:** 10.1101/2024.05.23.595560

**Authors:** Rajleen K. Gogar, Nidhi Chhikara, Minh Vo, Nathaniel C. Gilbert, Jack A. Dunkle, Patrick A. Frantom

## Abstract

Fe-S clusters are critical cofactors for redox chemistry in all organisms. The cysteine desulfurase, SufS, provides sulfur in the SUF Fe-S cluster bioassembly pathway. SufS is a dimeric, PLP-dependent enzyme that uses cysteine as a substrate to generate alanine and a covalent persulfide on an active site cysteine residue. SufS enzymes are activated by an accessory transpersulfurase protein, either SufE or SufU depending on the organism, which accepts the persulfide product and delivers it to downstream partners for Fe-S assembly. Here, using *E. coli* proteins, we present the first X-ray crystal structure of a SufS/SufE complex. There is a 1:1 stoichiometry with each monomeric unit of the EcSufS dimer bound to one EcSufE subunit, though one EcSufE is rotated ∼7° closer to the EcSufS active site. EcSufE makes clear interactions with the α16 helix of EcSufS and site-directed mutants of several α16 residues were deficient in EcSufE binding. Analysis of the EcSufE structure showed a loss of electron density at the EcSufS/EcSufE interface for a flexible loop containing the highly conserved residue R119. An R119A EcSufE variant binds EcSufS but is not active in cysteine desulfurase assays and fails to support Fe-S cluster bioassembly in vivo. ^35^S-transfer assays suggest that R119A EcSufE can receive a persulfide, suggesting the residue may function in a release mechanism. The structure of the EcSufS/EcSufE complex allows for comparison with other cysteine desulfurases to understand mechanisms of protected persulfide transfer across protein interfaces.

## Introduction

Cysteine desulfurases are PLP-dependent enzymes that catalyze the breakage of the C-S bond in cysteine to form alanine and a covalent cysteine persulfide species. These enzymes have been found in all three domains of life and are essential for the mobilization of sulfur for the biosynthesis of iron-sulfur clusters, thiol-containing cofactors, and thiolation modifications to tRNA molecules.(1) Cysteine desulfurases share an overall dimeric aminotransferase-V fold with two active sites made up of residues from both monomers (Figure 1A). Several active site residues are universally conserved including a lysine to form the internal PLP-aldimine, an active site cysteine to act as a nucleophile in C-S bond breakage step, and an arginine used to orient the carboxylate of the PLP-cys aldimine species for deprotonation of the α-proton. The shared active site architecture suggests that cysteine desulfurases utilize a common mechanism with a 1,3-proton shift of the PLP-cys aldimine to generate a PLP-cys ketimine intermediate prior to C-S bond cleavage.

**Figure 1.**
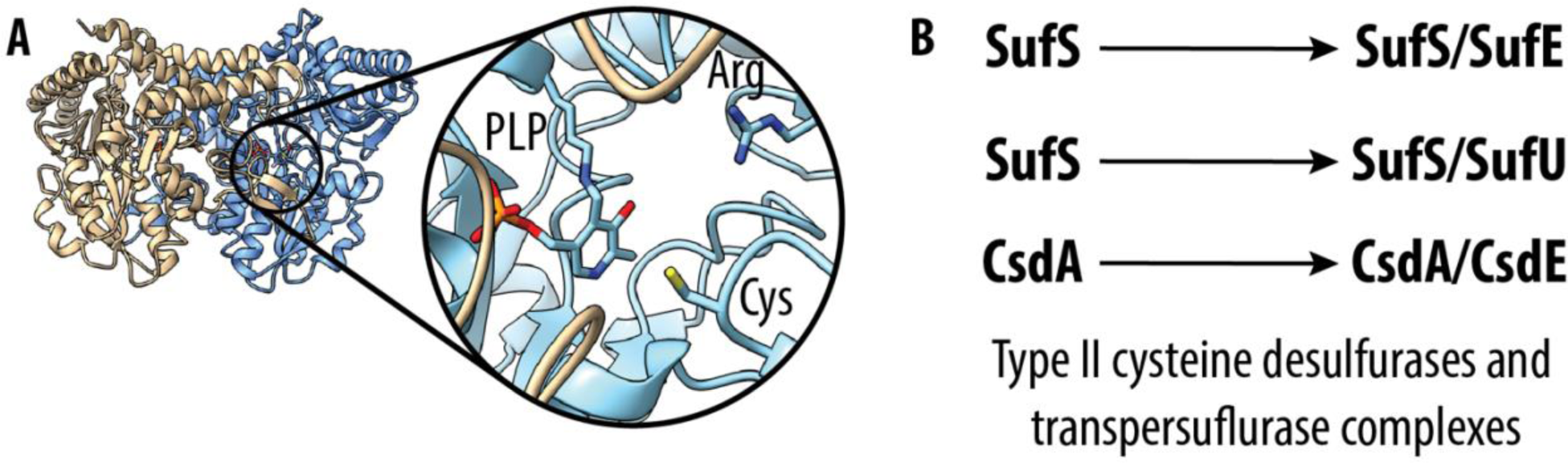
Overview of SufS and persuflide transfer. (A) Dimeric structure of EcSufS (PDB 6mr2). EcSufS monomers are colored tan and blue. The inset shows a magnified view of the active site with the PLP and active site cysteine residue. (B) List of homologous type II cysteine desulfurase/transpersulfurase pairs.

Despite the similarities, closer analysis of cysteine desulfurase structure and function reveals two distinct clusters of this large family of enzymes. Type I cysteine desulfurases, such as IscS and Nfs1, are found in bacteria and eukaryotic mitochondria.(2, 3) They can be identified structurally by an insertion near the active site cysteine, which places this key residue on a long, flexible loop. It is thought that the flexible S-transfer loop allows for persulfide transfer to a variety of acceptor proteins. Type II cysteine desulfurases are best represented by SufS in bacteria and archaea.(4) The active site cysteine residue in type II systems is found on a very short, stable S-transfer loop buried in the active site. Due to the buried nature of the active site, these enzymes require a dedicated accessory protein called a transpersulfurase to act as a second substrate in the reaction. The transpersulfurases contain an active site cysteine on an S-transfer loop that can extend into the desulfurase active site and accept the persulfide to complete the catalytic cycle.(5–7) The presence of the cognate transpersulfurase can accelerate the rate of SufS enzymes by 10-100 fold by acting as a preferred persulfide acceptor. Some of the type II systems have the additional attribute of providing protected persulfide transfer under oxidative stress conditions adding mechanistic importance to the desulfurase/transpersulfurase complex.(8, 9)

A survey of type II systems identifies three major complex pairs: SufS/SufU, SufS/SufE, and CsdA/CsdE (Figure 1B). SufS catalyzes the first step in the biosynthesis of iron-sulfur clusters in the SUF pathway found in bacteria, plants, and archaea.(4) In the Suf system, the identity of the transpersulfurase is organism specific with SufS/SufU complexes found in gram-positive and gram-negative bacteria and SufS/SufE favored in gram-negative bacteria and plants. The CsdA/CsdE system is confined to a narrow taxonomic range compared to the two Suf systems. The function of CsdA/CsdE is less clear but has been shown to be required for cyclization of the *N*^6^-threonylcarbamoyladenosine tRNA modification in *E. coli*, though the exact mechanistic contribution is currently unknown.(*10*)

Structures of SufU-containing complexes have been reported from multiple sources including *Bacillus subtilis*(*11*), *Mycobacterium tuberculosis*(*9*), and *Staphylococcus aureus*(*12*). Analysis of the structures along with complementary biochemical results indicates that the SufS/SufU systems require divalent zinc ions for activity. The zinc ion is initially coordinated within SufU with the acceptor cysteine residue as a ligand. Formation of the SufS/SufU complex results in a metal-ligand exchange reaction releasing the acceptor cysteine from SufU and mediating the SufS/SufU interface.(11) Prior to this report, there were no structures of a SufS/SufE complex, and structural representation of an E-type transpersulfurase complex was exclusively found in co-crystals of the CsdA-CsdE complex from *E. coli*.(13) However, there are several mechanistic and structural differences between the CsdA/CsdE system and the SufS/SufE system that suggest it is not a great mimic of the persulfide transfer mechanism in the Suf system. The active site of CsdA is more exposed than SufS due to a 50° bend in the α4 helix covering the active site.(13) This open active site is likely responsible for the fact that CsdE interacts with CsdA via a single binding step with high affinity(14) and only activates the desulfurase activity two-fold(15). In contrast, the SufS/SufE complex is proposed to use a two-step binding model(16), to activate SufS desulfurase activity to a much greater degree(17), and to protect persulfide transfer from exogenous oxidants and reductants(8). Lack of an iron-sulfur cluster specific SufS/SufE structure presents numerous challenges to understanding the mechanism of protected persulfide transfer in this pathway.

Here, we report the first co-structure for the SufS-SufE complex from *E. coli*. The overall structure contains one EcSufE monomer per EcSufS active site and is asymmetric with one EcSufE positioned closer to the EcSufS active site, though neither pairing is in a persulfide transfer ready position. A conserved arginine residue (R56) on the α3-α4 loop of EcSufS appears to be blocking EcSufE from making a closer approach. Additionally, both EcSufE subunits are missing density for a loop containing the strictly conserved R119. The structural results are complemented by site-directed mutagenesis studies that identify several residues on the α16 helix of EcSufS as essential for initial EcSufE binding. Several of these residues are shared among the type II cysteine desulfurases with the exception of Y345, suggesting it is specific to SufS/SufE interactions. Investigations of residues on the R119-containing mobile loop identified R119 as an essential residue for catalytic turnover of the EcSufS/EcSufE system. Radiolabel transfer assays show that R119A EcSufE can accept a persulfide from EcSufS pointing towards a role for R119 in breaking the EcSufS/EcSufE complex following persulfide transfer. The details gleaned from the data presented here are compared with the SufS/SufU and CsdA/CsdE complexes to generate a proposed molecular mechanism for protected persulfide transfer in SufS/SufE catalysis.

## Results

### Structural characterization of the EcSufS/EcSufE complex

In a previous report, we created a double variant of EcSufE (C51A/E107C) lacking the active site C51 residue.(16) This variant was suitable for labeling with a maleimide-BODIPY FL fluorescent tag to use as a binding probe in fluorescence polarization assays with EcSufS. In this series of experiments, we were able to demonstrate that addition of cysteine substrate resulted in a 10-fold improvement in the *K*_D_ value for formation of the EcSufS/EcSufE complex (from ∼5 µM to ∼0.5 µM). This result could be explained by loss of the persulfide-accepting C51 residue in C51A/E107C EcSufE leading to a trapped, high-affinity EcSufS-S^−^/EcSufE intermediate. We hypothesized that this intermediate could be used for crystallography trials in an attempt to capture an EcSufS/EcSufE complex. Solved crystal structures from this experiment resulted in a EcSufS/EcSufE complex with 1:1 stoichiometry between the S and E subunits (Figure 2 and Table S1).

**Figure 2.**
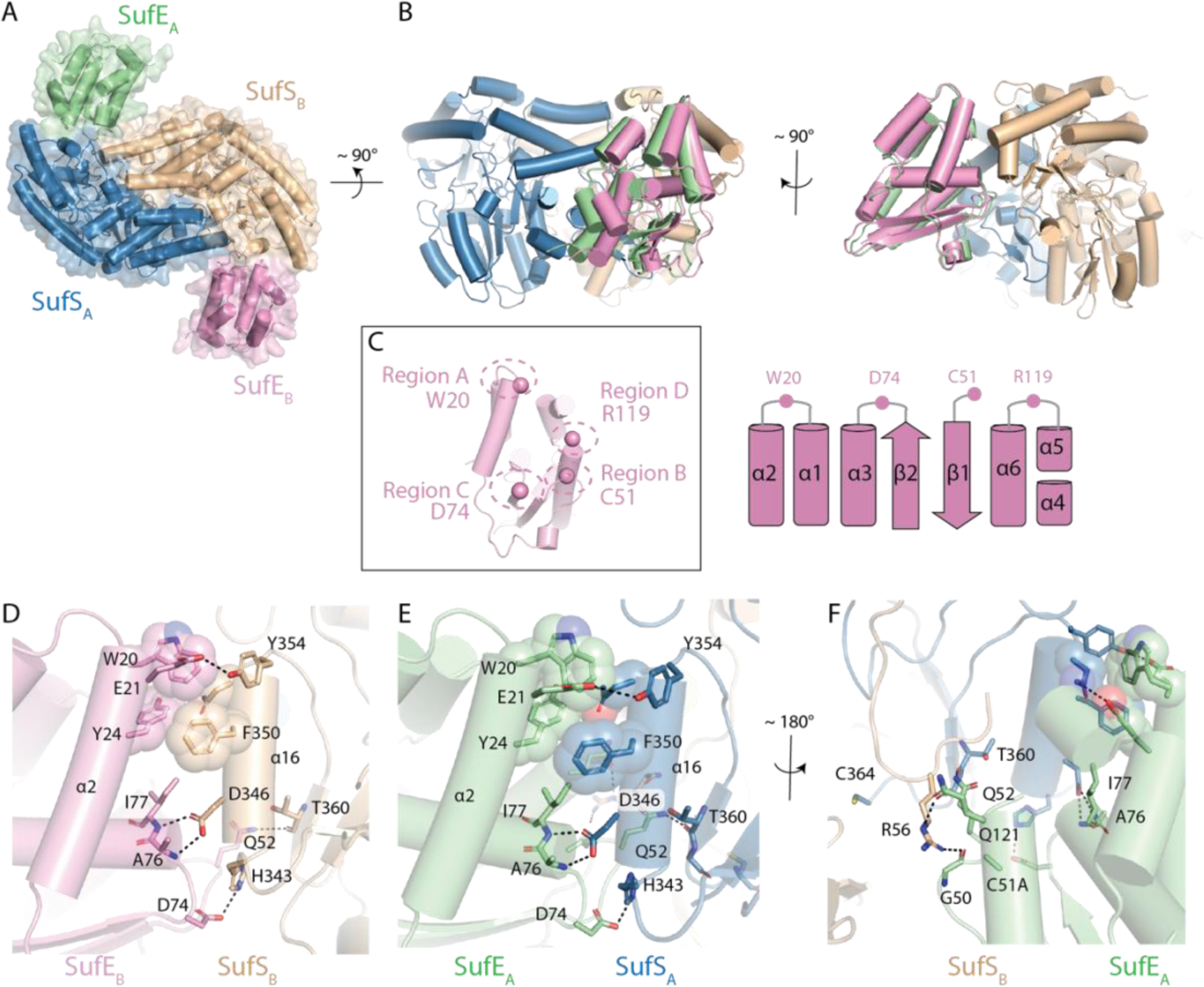
Four regions of SufE form contacts to SufS. (A) The X-ray crystal structure of the SufS homodimer bound to two copies of SufE. (B). The two copies of SufE are not superimposable with SufE_A_ (green) rotated more towards SufS, forming more intimate interactions with SufS than its SufE_B_ (pink) counterpart. (C) Four regions of SufE contact SufS, labeled regions A to D. A critical amino acid within each of these regions is given in one-letter code, W20, C51, D74 and R119. A topology diagram shows where each of the four regions is located within SufE. (D) The detailed interactions of SufE_B_ with helix α16 of SufS_B_. (E)The detailed interactions of SufE_A_ with helix α16 of SufS_A_ which are very similar to those of SufE_B_ with SufS_B_. (F) SufE_A_ forms additional interactions with R56 of SufS_B_ not seen in SufE_B_ that explains it closer approach the SufS homodimer.

The positioning of the EcSufE subunits creates asymmetry in the overall structure, but superposition of the two EcSufS monomers shows no major changes in Cα backbone or in the side chain orientations for key regions. The same is true for the isolated EcSufE subunits. The overall asymmetry is best described as a rigid rotation of EcSufE towards EcSufS resulting in a closure of approximately 7° (Figures 2A and 2B) The more open interaction is based primarily on contacts between EcSufE and the α16 helix (residues 343-353 in EcSufS) of EcSufS (Figure 2D and 2E) while the closure creates additional interactions between the C51 and R119 loops on EcSufE with the R56 loop and α4 helix of the adjacent SufS monomer (Figure 2F). In both cases, the C51 loop of EcSufE remains in an inward facing position.

Although this is the first reported structure of a SufS/SufE complex, homologous structures of the CsdA/CsdE complex from *E. coli* and the SufS/SufU complex from several organisms identified the α16 helix (residues 343-353 in EcSufS) as a primary point of contact between the two proteins. Prominent hydrogen bonding contacts occur between EcSufE and three residues present on α16 of EcSufS (Figure 2E). H343, D346, and Y354 of EcSufS hydrogen bond to D74, backbone amides of A76 and I77, and E21 on EcSufE, respectively. Two of these interactions have previously been investigated. The D346R substitution in EcSufS results in loss of interaction with EcSufE(16), while the D74R EcSufE variant paradoxically binds to EcSufS with a 10-fold improvement in *K*_D_ value(18). The remainder of the α16 helix interface is made up of non-polar interactions between EcSufE and Y345 and F350 on EcSufS. These interactions are similar to those seen in the previously reported CsdA/CsdE structure but are not conserved in the reported SufS/SufU complex structures.

The more closed EcSufS/EcSufE pairing creates additional contacts between EcSufE and the adjacent EcSufS monomer. The EcSufE residues in these contacts are located in the C51 loop (residues 47-53) and the R119 loop and flanking regions (residues 112-128); however, the loop surrounding R119 (residues 116-120) lacks density in both EcSufE subunits (Figures 3A and 3B). These two regions form layered interactions with the N-terminus of the α4 helix (residues 60-67) and the adjacent R56 loop (residues 56-59) on EcSufS, with R56 appearing to block formation of a closer complex. These interactions are not seen in the CsdA/CsdE structure. In this case, the α4 helix and R56 loop undergo a dramatic conformational change and shift away from the interaction with CsdE.(13) In contrast, the interactions in this region are similar in architecture to those seen in the SufS/SufU complex (11).

**Figure 3.**
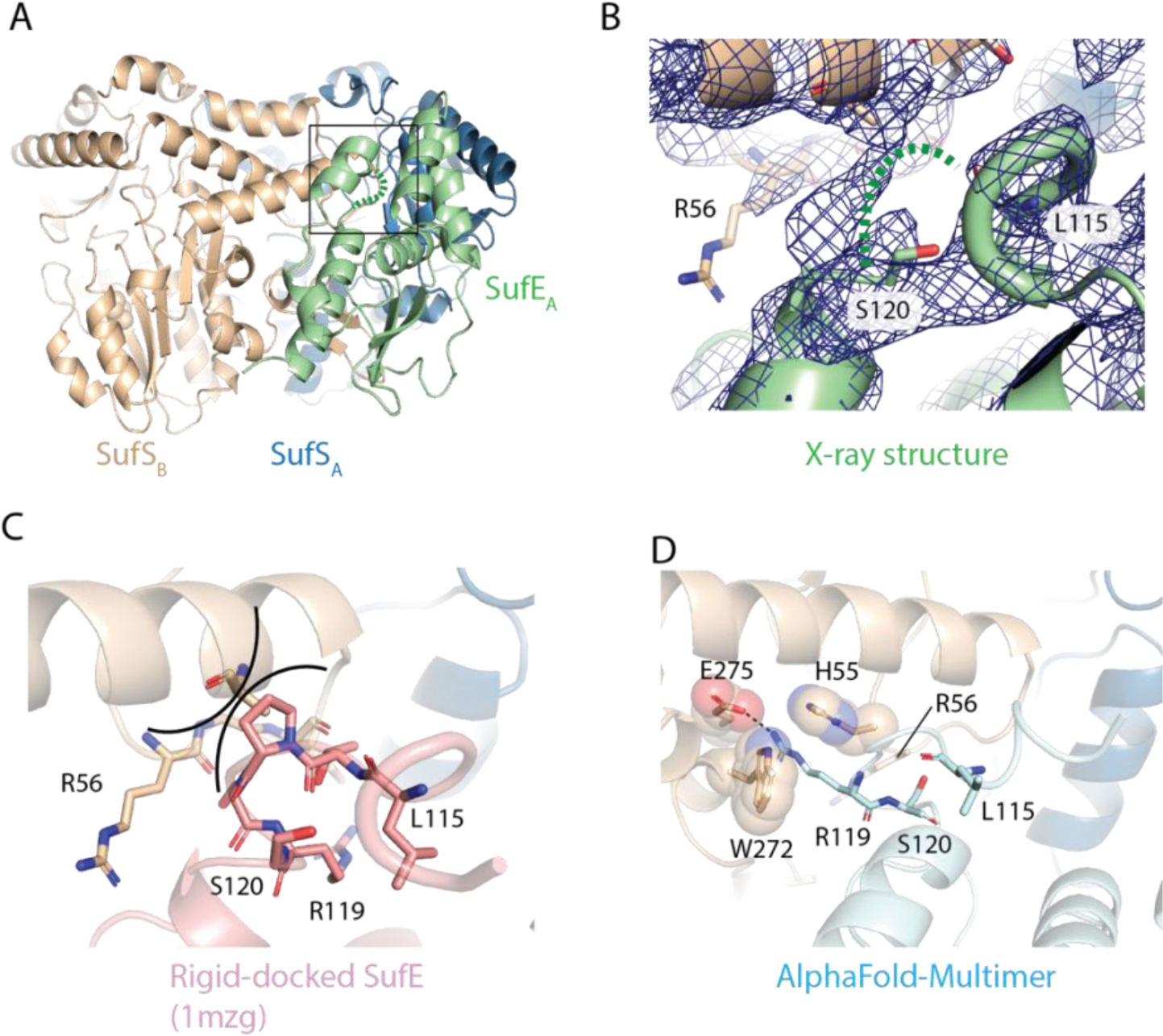
Conformational changes in the SufE R119 loop upon interaction with SufS. (A) The boxed area shows the location where the R119 loop of SufE interacts with SufS. (B) The SufSE X-ray structure lacks density for residues T116-R119 indicating a dynamic unfolding event upon interaction with SufE. A 2F_o_-F_c_ electron density map is shown (blue) contoured at 1.0 σ. (C) Rigid-docking of the isolated X-ray structure of SufE onto its position within the SufSE complex indicates steric clashes (indicated by arcs) with SufS will occur unless there is a conformational change in SufE. (D) An AlphaFold2-multimer model of the SufSE interaction predicts a conformational change of the loop with R119 forming cation-π and electrostatic interactions with W272, H55 and E275 respectively.

The loss of density for the R119 loop in both EcSufE monomers suggests that it is highly flexible in this version of the complex. A rigid-docked version of a EcSufE monomer based on PDB code 1mzg shows that some conformational change is required or the R119 loop would clash with residues on the α4 and α16 helix (Figure 3C). AlphaFold2 was used via ColabFold(19) to predict the location of the R119 loop in the EcSufS/EcSufE complex. The modeled subunits strongly resemble the experimental complex with an RMSD of 0.45 Å when aligned on an EcSufS monomer and 0.62 Å RMSD between the EcSufE subunits. The main difference in EcSufE structures is in the conformation of the R119 loop. In the AlphaFold2 model, the loop is extended and R119 enters a divot on EcSufS adjacent to the active site. In this conformation R119 forms an electrostatic interaction with E275 of EcSufS and hydrogen bonds to the backbone carbonyl of S254 (Figure 3D). While the modeled R119 loop is labeled as “low confidence” by AlphaFold2, experiments described below provide support for a mechanistic role for this residue.

### Biochemical analysis of EcSufS/EcSufE interactions in EcSufS α16 helix

The α16 helix of EcSufS appears to provide the initial binding face for EcSufE.(16) This helix consists of residues 343-355 in EcSufS. The outward facing residues that have contact with EcSufE include H343, Y345, D346, S349, F350, N353, and Y354 (Figure 4A). A survey of type II cysteine desulfurase sequences identifies D346 as the only strictly conserved residue. To determine the importance of each residue on the helix in the EcSufS/EcSufE interaction, alanine-scanning mutagenesis was used to generate single site variants of EcSufS. All variant proteins were expressed and purified similar to the wildtype protein (Figure S1), and results from circular dichroism spectroscopy suggest that none of the substitutions dramatically altered the structure of EcSufS (Figures S2). These variants were screened for the ability to bind to EcSufE via a fluorescence polarization assay (Table 1). S349A and N353A gave *K*_D_ values in the 0.1-2 µM range, similar to or better than values determined with the wildtype EcSufS (Figure 4B). Apparent *K*_D_ values of >20 µM were determined for three of the substitutions (H343A, Y345A, and Y354A) (Figure 4C). Substitution of D346 or F350 with alanine resulted in a complete loss of the EcSufS interaction with EcSufE at concentrations up to 20 µM (Figure 4D). When the effects on *K*_D_ value are mapped onto the EcSufS structure, co-localized patches emerge providing a clearer picture of the functional interactions between EcSufS and EcSufE (Figure 4A).

**Figure 4.**
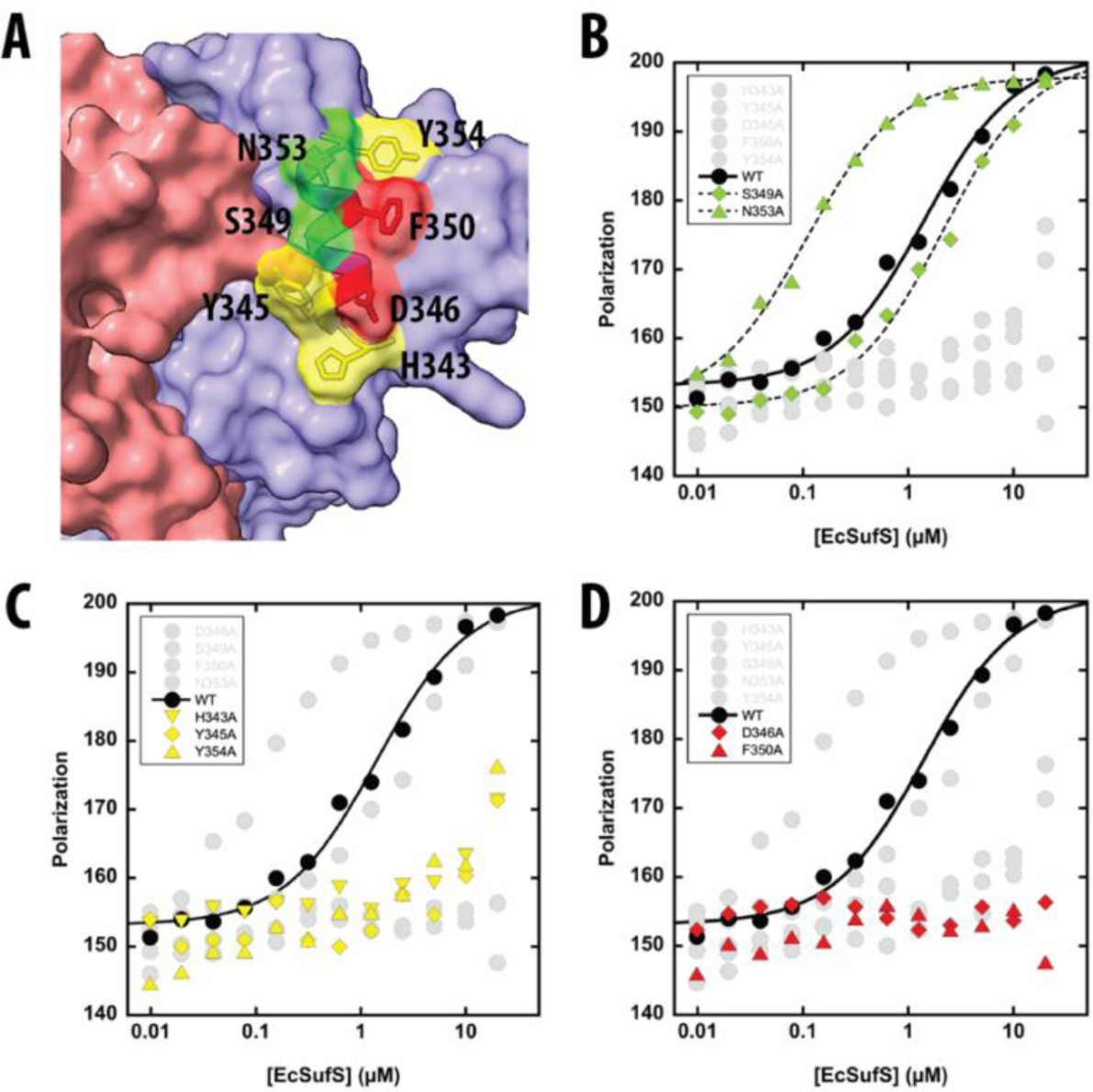
Role for residues on α16 in binding EcSufE. (A) Residues on the α16 helix investigated via site-directed mutagenesis and color coded based on the effect on EcSufE interactions. Representative fluorescence polarization data for EcSufS and variants displaying (B) strong interactions (green), (C) weak interactions (yellow), and (D) no interactions (red) with EcSufE. Full triplicate data sets are shown in Figure S4.

**Table 1.** Apparent binding constants for the EcSufS/EcSufE interaction measured by fluorescence polarization.^a^.

| enzyme<sup>b</sup> | $K_D$ in absence of cysteine ($\mu\text{M}$) | $K_D$ in presence of cysteine ($\mu\text{M}$) |
| --- | --- | --- |
| SufS/SufE | $5.2 \pm 1.5$ | $0.6 \pm 0.2$ |
| H343A SufS | >20 | N.D.<sup>c</sup> |
| Y345A SufS | >20 | N.D. |
| D346A SufS | >>40 | N.D. |
| S349A SufS | $2.2 \pm 0.7$ | N.D. |
| F350A SufS | >>40 | N.D. |
| N353A SufS | $0.10 \pm 0.03$ | N.D. |
| Y354A SufS | >20 | N.D. |
| T116A SufE | $2.3 \pm 0.6$ | $0.9 \pm 0.3$ |
| T116S SufE | $3.4 \pm 2.0$ | $3.3 \pm 2.0$ |
| R119A SufE | $0.4 \pm 0.1$ | $0.4 \pm 0.1$ |
| R119K SufE | $1.5 \pm 0.8$ | $0.9 \pm 0.4$ |
| Q121A SufE | $6.2 \pm 3.0$ | $3.1 \pm 1.1$ |
| Q121E SufE | $6.4 \pm 2.2$ | $0.7 \pm 0.9$ |

<sup>a</sup>Experimental conditions described in Methods section. <sup>b</sup>The variant enzymes listed were tested against their wildtype partner. <sup>c</sup>Not determined.

### Functional importance of the R119 EcSufE residue

Due to the lack of defined structure, the residues on the R119 EcSufE loop were investigated using site-directed mutagenesis. Three residues were selected based on conservation within SufE sequences and location in the AlphaFold2 model (Figure 5A). T116, R119, and Q121 were all substituted with alanine as well as a more conservative residue to generate six variants of EcSufE in both the wildtype and C51A/E107C backgrounds. SDS-PAGE (Figure S1) and circular dichroism spectra (Figure S2) confirm the purity of each EcSufE variant and the lack of perturbation to secondary structure. Substitution in the wildtype enzyme were used to determine Michaelis-Menten parameters for the EcSufE substrates in the presence of saturating cysteine. All of these enzyme variants exhibited kinetic parameters identical to the wildtype enzyme with the exception of the R119A EcSufE enzyme (Table 2). This variant did not display cysteine desulfurase activity at concentrations up to 3 µM EcSufE; however, the R119K substitution rescued desulfurase activity (Figure 5B). Identical substitutions made in the C51A/E107C EcSufE enzyme were labeled with BODIPY-FL-maleimide and used in the fluorescence polarization assay to determine *K*_D_ values for the EcSufS/EcSufE interaction (Table 1). Due to the background C51A mutation in EcSufE used in this experiment, the *K*_D_ values can be determined in the absence (Figure S4) or presence of cysteine (Figure S5). We have previously reported that the addition of cysteine improves the *K*_D_ value from ∼5 µM to ∼0.5 µM, likely due to the formation of the EcSufS active site persulfide and a more realistic persulfide transfer complex.(16) For the R119-loop variants, the *K*_D_ values determined in the absence or presence of cysteine were not largely different that the values determined with the C51A/E107C EcSufE control, with none of the substitutions resulting in large binding defects. Surprisingly, the R119A substitution exhibited the lowest *K*_D_ values despite not being active in the cysteine desulfurase assay (Figure 5C). This result suggests that the lack of catalytic activity with the R119A EcSufE is not due to a lack of binding.

**Figure 5.**
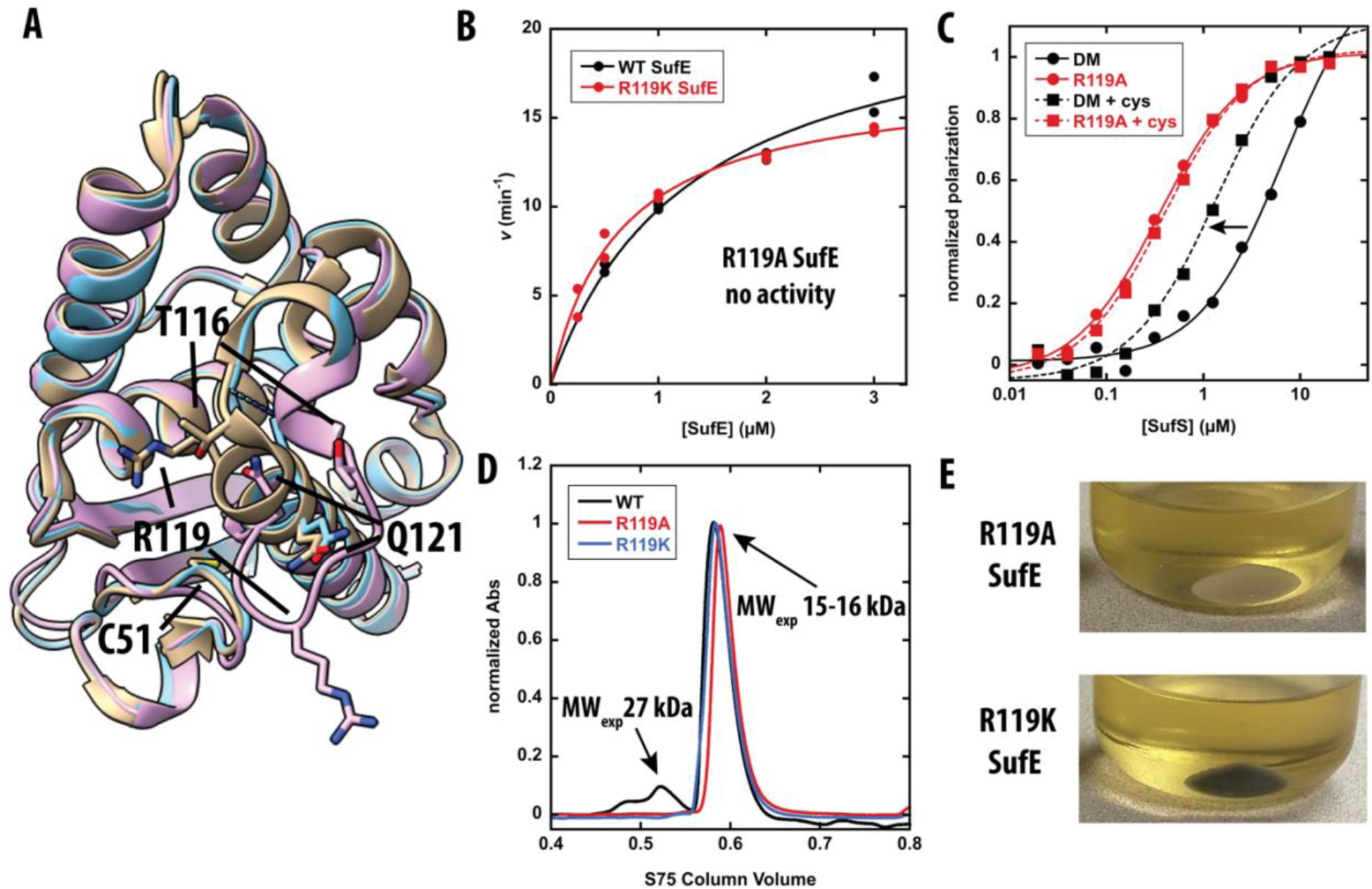
Biochemical characterization of EcSufE R119 residue. (A) Superposition of EcSufE structures from unbound (tan, 1mzg), in complex with EcSufS (blue, 8vbs), and the AlphaFold2 model (pink). (B) Michaelis-Menten curve for alanine production by wildtype (black) and R119K (red) EcSufE. Lines are from fits of the data to equation 1. (C) Fluorescence polarization assay results for “wildtype” EcSufE (black, labeled as Double Mutant for background C51A/E107C mutations) and R119A EcSufE (red, in DM background) in the absence (circle, solid line) and presence (square, dashed line) of 500 µM cysteine. Lines are from fits of the data to equation 2. The arrow denotes the improvement in K_D_ value for the “wildtype” EcSufE after addition of cysteine. (D) S75 analytical size exclusion chromatography chromatograms for wildtype EcSufE (black), R119A EcSufE (red), and R119K EcSufE (blue). Molecular weights were estimated as described in Materials and Methods. (E) Cell pellets for expression of the *suf* operon in *E. coli* cells. Black cell pellets indicate a functional Fe-S cluster assembly pathway.

**Table 2.**
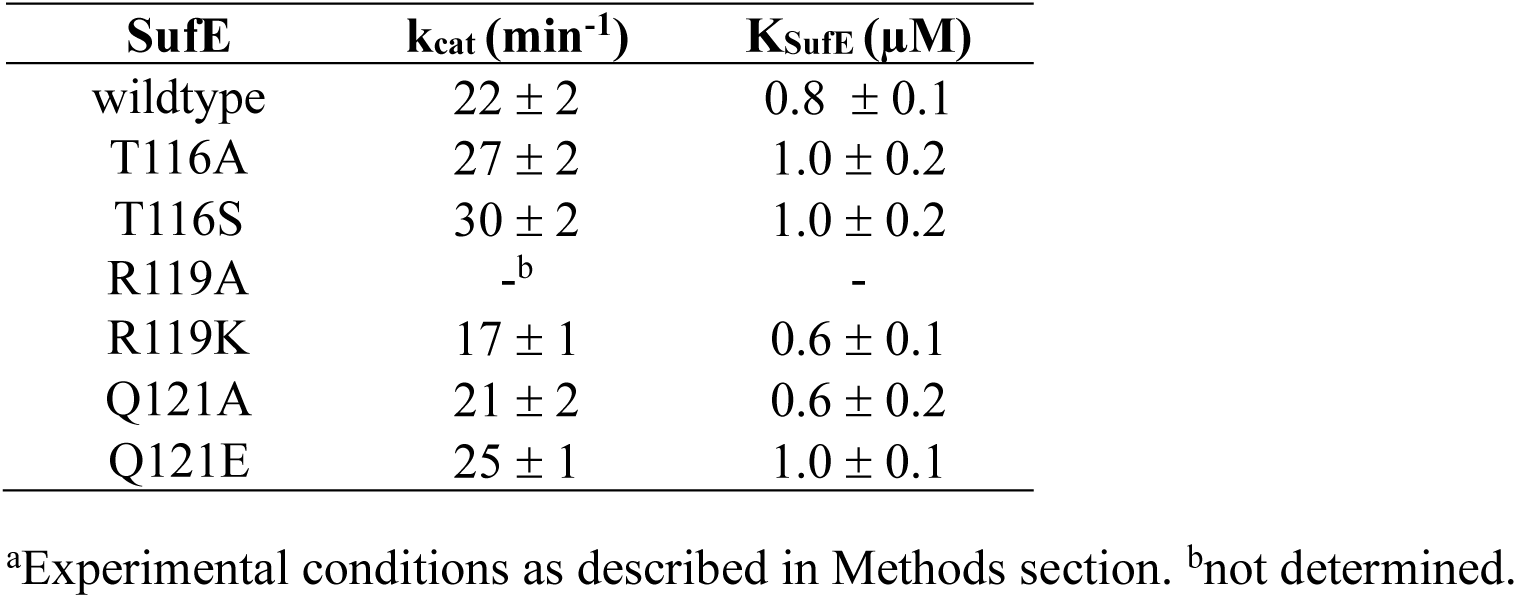
Michaelis-Menten parameters for EcSufE R119 loop variants^a^.

Several additional experiments were performed to confirm the essential nature of the R119 EcSufE residue. In the EcSufE structure reported in PDB code 1mzg, the protein crystallizes as a dimer with R119 forming an electrostatic interaction with E21’ from the adjacent monomer (Figure S6). EcSufE has previously been shown to primarily form a monomer in-solution, and E21 is not well-conserved suggesting the dimeric form is not critical to EcSufE function. To ensure that the R119A substitution had not changed to quaternary structure of EcSuE, size-exclusion chromatography was used to estimate a native molecular weight for wildtype, R119A, and R119K EcSufE (Figure 5D). The data shows that all three proteins elute from an S75 column with an estimated molecular weight of 15 kDa, consistent with a monomeric form. Finally, to determine if the R119A substitution was sufficient to affect Fe-S cluster formation in vivo, the same mutation was made in a pDuet plasmid that carries the SUF operon (*suf*ABCDSE). Expression of the functional operon in *E. coli* results in production of Fe-S clusters and a dark black cell pellet. When the R119A substituted operon is induced under identical conditions, the cell pellet is pale brown (Figure 5E). Expression of the R119K-containing plasmid rescues the Fe-S cluster assembly phenotype. Overall, these results are consistent with the R119 EcSufE residue as catalytically necessary for EcSufS cysteine desulfurase activity and Fe-S cluster assembly in vivo.

### 35S transfer assay shows R119A EcSufE can accept the persulfide from EcSufS

Based on the previous results, two mechanisms emerge to describe the R119A EcSufE defect. One possibility is that R119A EcSufE forms the “close-approach” complex with EcSufS and inhibits the reaction by being unable to accept the persulfide. A second possibility is that R119A EcSufE is capable of accepting the persulfide from EcSufS but is not able to escape the “close-approach” conformation, effectively inhibiting the reaction. To discriminate between these two mechanisms, ^35^S-cysteine was incubated with various forms of EcSufS and EcSufE in the absence of reductants. SDS-PAGE was then used to separate the proteins, and the gel was imaged with a phosphoimager to detect ^35^S incorporation in the form of persulfide (Figure 6). In the presence of EcSufE, R119A EcSufS (lane 5) is labeled with ^35^S to a similar extent as the wildtype EcSufE (lane 3) and the fully functional R119K EcSufE (lane 7). This result supports a mechanism of R119A EcSufE inhibition based on an inability to escape the “close-approach” complex.

**Figure 6.**
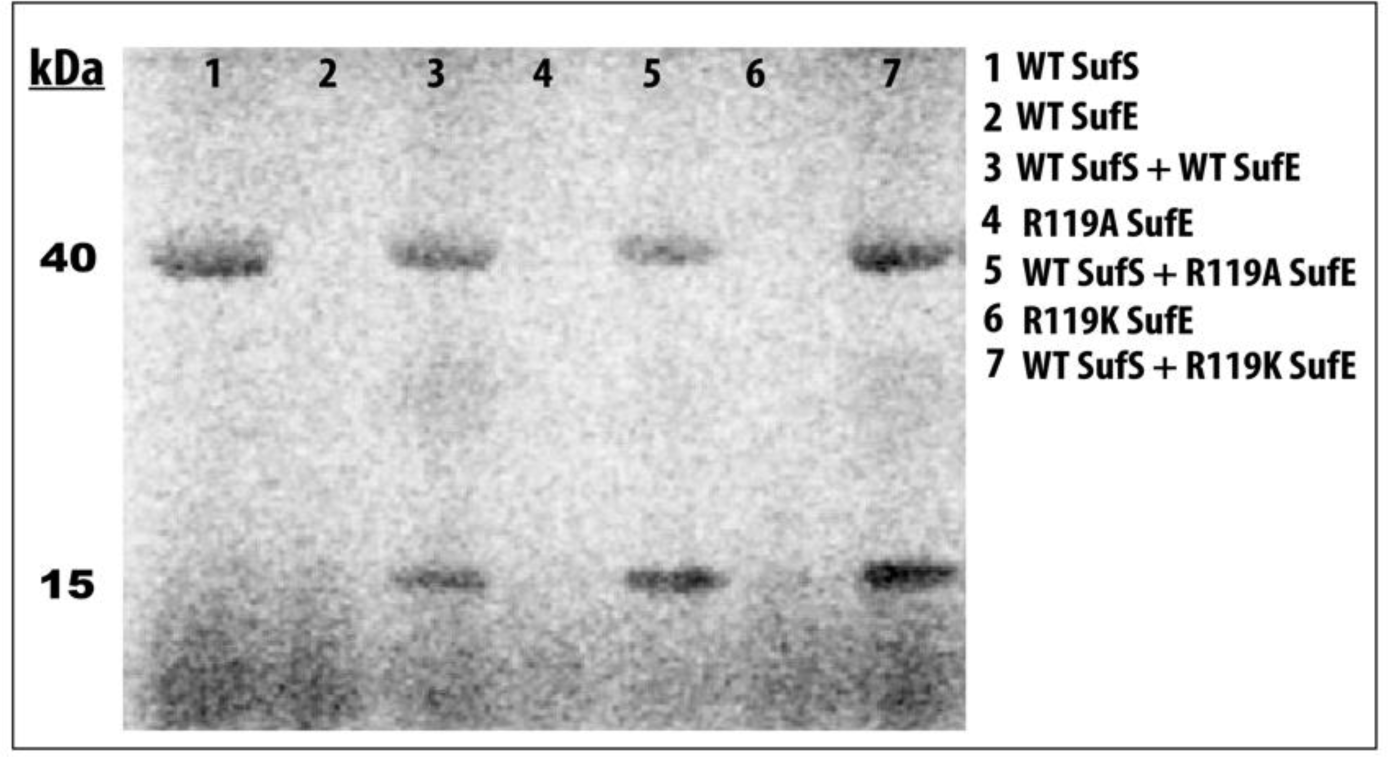
Persulfide formation reaction with ^35^S-cysteine monitored by SDS-PAGE. Reactions were run as described in the methods section. Labels indicate the identities of SufS (44 kDa) or SufE (15 kDa) used in each reaction.

### Sequence similarity network for SufE proteins reveals conserved regions of the fold

A sequence similarity network was created for the SufE-containing InterPro family IPR003808 containing 1447 representative nodes dispersed into monotaxonomic clusters (Figure 7A). To get a clear picture of the diversity of sequences contributing to the SufE fold and obtain unbiased estimates of position-specific conservation, three sequences from each major cluster were selected and used to generate a multiple sequence alignment (Figure S7). Based on the alignment, hidden-Markov model (HMM) logos were generated for areas of interest. The HMM logos in Figure 7B were created to visualize the sequence conservation at the SufS/SufE interface and show two universally conserved residues across the SufE-like folds. One is the active site cysteine residue (C51 in EcSufE) in Region B, which is required for transpersulfurase activity. The other is an arginine residue (R119 in EcSufE) in Region D. Indeed, these two residues are among a very small number of strictly conserved residues in the SufE-like folds (Figure S7).

**Figure 7.**
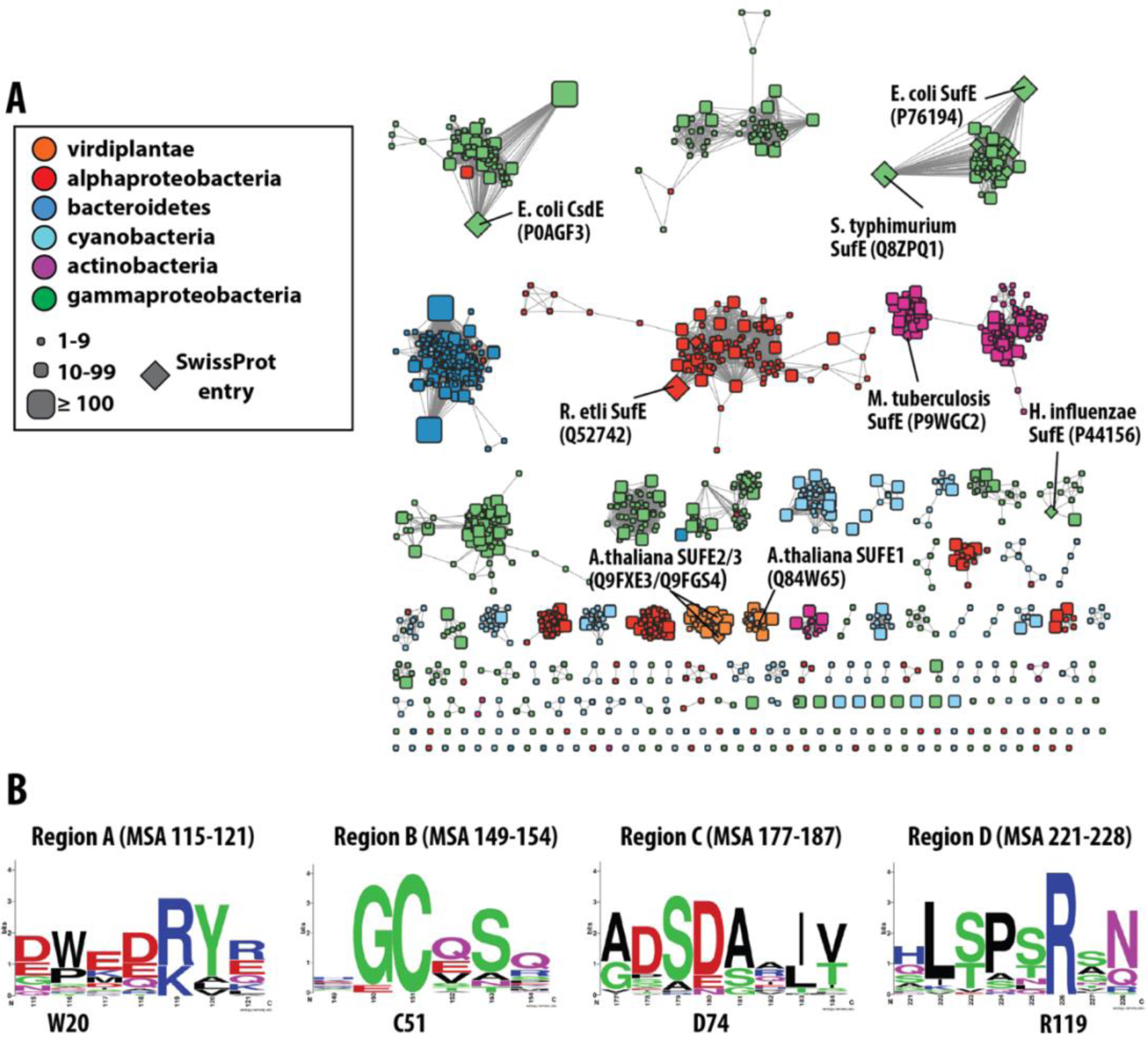
Sequence similarity network of the SufE-like domain (IPR003808). (A) A representative sequence similarity network for the IPR003808 domain visualized at an alignment cutoff value of 45. (B). HMM logos generated from an unbiased multiple sequence alignment of representative sequences from the network. Region labels correspond to EcSufS/EcSufE interaction sites outlined in Figure 2C.

## Discussion

### The EcSufS/EcSufE structure reveals interfacial residues required for complex formation and suggests a mechanism for transpersulfurase loop opening

Cysteine desulfurases are a major pathway for sulfur metabolism in living systems. Instead of free sulfur, these enzymes produce a covalent persufide product that must be transferred for catalytic turnover. While cysteine desulfurase mechanisms to generate the persulfide are strongly conserved, the fate of the persulfide varies based on the acceptor. Here, we report the first structure of a SufS/SufE cysteine desulfurase/transpersulferase complex. This complex is required for Fe-S cluster formation in the SUF pathway and protects the persulfide intermediate from external reductants and oxidants. A comparison of SufS homologs (SufS/SufE, SufS/SufU, and CsdA/E) with their various transpersulfurase partners shows conserved residues on the α16 helix with an HX**<u>X</u>**DXXζΦ motif, where **<u>X</u>** indicates a specificity residue, ζ indicates a hydrophilic residue, and Φ indicates a hydrophobic residue (Figure 8A). Binding assays described above confirm that the conserved H343, D346, and F350 (Φ) are important residues for forming an initial complex between EcSufS and EcSufE based on defects in *K*_D_ values for complex formation (Figure 3). Their conservation across SufS homologs with different transpersulfurase partners suggests they may play similar structural roles in complex recognition (Figure 8B-D). However, the similarity at these positions is not sufficient to promote cross-reactivity between systems, implicating additional features that drive specificity.

**Figure 8.**
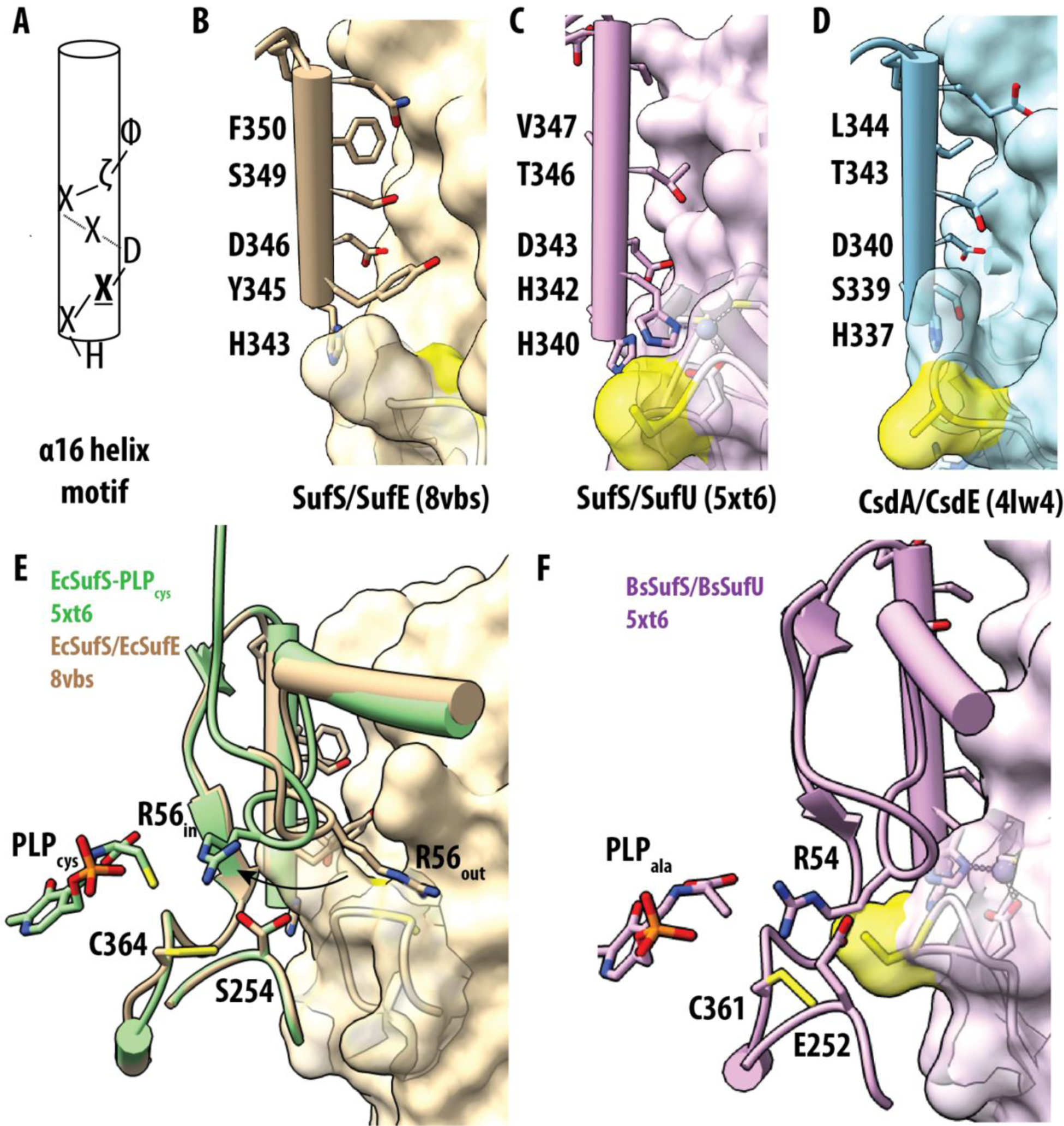
Structural comparison of type II cysteine desulfurases at the transpersulfurase interface. (A) Consensus figure for the α16 helix sequence. B-D) Structural representation of the α16 helix residues and the partner transpersulfurase for EcSufS/EcSufE (tan, B), BsSufS/BsSufU (purple, C), and EcCsdA/EcCsdE (blue, D). The residue containing the transpersulfurase active site cysteine is shown in yellow. (E) Structural representations changes in the α3-α4 loop from the EcSufS/EcSufE structure (tan) to those seen in the catalytic intermediate PLP-cys aldimine structure (green). (F) Structural representations of the α3-α4 loop and partner transpersulfurase for BsSufS/BsSufU to highlight similarities in arginine placement.

The **<u>X</u>** residue is unique to each SufS homolog and appears to play an important role in specificity and possibly opening the transpersulfurase loop. This is best characterized in the SufS/SufU systems where a histidine residue on SufS participates in a metal-ligand exchange between the persulfide-accepting cysteine residue and a divalent Zn^2+^ ion coordinated to SufU. (9, 11) The ligand exchange allows the SufU S-transfer loop to enter the SufS active site and promotes persulfide transfer (H342 in Figure 8C). In the EcSufS/EcSufE system, this residue is a tyrosine (Figure 8A). In the rigid docked model of EcSufS/EcSufE, Y345 forms a direct conflict with the R119 residue on EcSufE (Figure 3C). This clash would necessitate the rearrangement of the R119 region on EcSufE consistent with the loss of density in the structure. Given the mechanistic role of the homologous histidine in SufS/SufU pairs, it is possible that this tyrosine could insert into the empty hydrophobic pocket to promote formation of the open complex of the C51 loop in EcSufE. In CsdA/CsdE systems, this residue is a serine residue, which makes interactions with the open loop structure of CsdE (S339 in Figure 8D).

### Formation of a “close-approach” complex and persulfide transfer may rely on a conserved arginine residue in the SufS α3-α4 mobile loop

Multiple binding studies on formation of the EcSufS/EcSufE complex support a two-step binding process with formation of an initial complex with a *K*_D_ value of ∼5-10 µM followed by formation of a “close-approach” complex with a *K*_D_ value < 1µM stimulated by activated versions of EcSufS or EcSufE.(16, 18, 20) However, in crystal structures of SufS homologs, the two cysteine residues (usually one -C-SH and one C-S-S^−^) involved in persulfide transfer loops are still too distant from each other for efficient transfer.(9, 11) This suggests an additional set of conformational changes is necessary for catalytic function. In our EcSufS/EcSufE complex, R56 on the α3-α4 mobile loop appears to block EcSufE from making a closer interaction with EcSufS (Figure 3C). In other EcSufS structures, R56 can be found in both “out” and “in” positions (Figure 8E). In the “out” position, R56 does not make clear contacts with any other residues. In the “in” position, it forms multiple interactions near the SufS active site and would not prevent approach of the transpersulfurase partner. Importantly, movement of the R56 loop may able be linked to SufS catalysis. We recently reported the structure of the EcSufS PLP-cys aldimine intermediate.(21) In this structure, the R56 residue contacts the sulfur atom on the PLP-cys intermediate (Figure 8E). Using R56 and the α3-α4 mobile loop to jointly coordinate SufS chemistry and SufE interactions is an attractive mechanism to regulate persulfide transfer. In the SufS/SufU systems, the analogous arginine residue is found in the “in” position in an electrostatic interaction with a glutamate residue (Figure 8F). This glutamate residue is in an analogous position to S254 in EcSufS. This residue lies in the middle of the recently identified β-latch motif, which also participates in formation of the close-approach conformation.(16)

### R119 is a critical residue for release of EcSufE from the “close-approach” conformation

The R119 loop in Region D of EcSufE lacked electron density at both interfaces suggesting that this region may be dynamic in nature (Figure 3B). As described above, rigid docking of the PDB 1mzg structure to the EcSufE subunit shows the loop will clash with both the α4 helix and with Y345 of EcSufE, the specificty **<u>X</u>** residue described above, necessitating a conformational change for this region in the complex (Figure 3C). Analysis of the sequence similarity network for the SufE fold highlights the conservation of the R119 residue suggesting it plays a functional role in persulfide transfer.

Results from fluorescence polarization assays are consistent with the R119A EcSufE variant being able to bind to EcSufS in an approximation of the “close approach” conformation, based on sub-micromolar *K*_D_ values, but unable to support catalytic turnover (Figure 4). This appearance of sub-micromolar dissociation constants mirrors the tight-binding nature of other activated EcSufE forms including the C51-alkylated form(22) and the D74R variant(18). In those cases, it is thought that the C51-loop is stabilized in the outward position. In the case of the R119A substitution, the current structure suggests that the C51 loop remains in the inward facing position and that R119 is not solely responsible for maintaining that position. Various control experiments confirm that changes in the function of the R119A variant are not the results of gross structural changes, and the R119K variant is able to rescue the defects created by the alanine substitution. The key result in establishing a mechanistic role for the R119 residue is the ability of R119A EcSufE to accept persulfide from EcSufS (Figure 7). The ability to accept persulfide suggests that the C51-loop is capable of forming a productive conformation in the absence of R119. Taken together, this set of results supports a mechanism where the R119 residue is essential for EcSufE to escape the “close approach” complex with EcSufS.

Structural comparison of the rigid docked EcSufS with the complex structure points toward a role for Y345 in the **<u>X</u>** position helping to open the EcSufE C51 loop to allow for a “close approach” conformation that necessitates movement of the R119 loop away from the interface. Indeed, the analogous arginine-containing loop in the CsdA/CsdE complex is rotated away from the CsdA interface, providing further evidence for the flexibility of this region.(13) SufU also contains an analogous arginine residue that forms a salt bridge with the aspartic acid from the HX**<u>X</u>**DXXζΦ SufS motif. This interaction is not perturbed by the histidine in the **<u>X</u>** position of SufS enzymes with SufU partners, which is involved in ligand metal exchange mechanism to open the SufU S-transfer loop. It is highly likely SufS/SufU complexes do not rely on the arginine for release and instead uses a second metal-ligand exchange allowing the S-transfer loop to move back to the inward facing position.

### A proposed mechanism for protected persulfide transfer in SufS/SufE systems

The requirement of SufE as a requirement for robust SufS activity was identified over 20 years ago.(5, 6) Although we do not have discrete crystallographic evidence for each step, reasonable evidence exists to propose a detailed molecular mechanism for persulfide transfer in the SufS/SufE systems that includes mechanisms for both inducing tight binding to promote persulfide transfer and overcoming tight binding to release SufE. Binding of SufE to SufS occurs through a two-step mechanism with a weak interaction (*K*_D_ ∼5-10 µM) focused around interactions with residues on the SufS α16 helix and requiring a conformational change of the R119-loop on SufE. Positioning of R56 on the α3-α4 mobile loop of SufS and a closed S-transfer loop on SufE prevent formation of the “close-approach” complex. Upon reaction of cysteine with the PLP of SufS, R56 moves to an inward position where it can coordinate with the sulfur atom of either the PLP-cys intermediate or the C364 persulfide as seen in the SufS PLP-cys aldimine structure and in SufS/SufU structures. This movement allows the C51 loop to move close enough to form cross-protein interactions with SufS leading to loop opening. As SufE moves toward persulfide acceptance, Y345 on SufS could insert into the cavity vacated by the C51 loop and force R119 into an outward facing conformation to avoid steric clashes. After persulfide transfer to SufE, the C51 persulfide loop closure requires R119 to return to its position to displace Y345 and allow the persulfide loop to fully close, re-creating the low-affinity complex. This step is supported by the ability of the R119A EcSufE variant to bind tightly in the absence of cysteine, accept persulfide from EcSufS, and the inability of the complex to support turnover.

## Conclusions

The protected persulfide transfer of the SufS/SufE system is essential to the function of the SUF pathway. Here, several structural features have been identified that provide a molecular mechanism for this activity. A flexible loop containing the conserved R119 on EcSufE must change conformation to allow for formation of an initial complex with residues on the α16 helix of EcSufS while the R56-containing α3-α4 loop of EcSufS is required to move inward to create a “close-approach” complex. Based on previous structural changes, it appears movement of the α_3_-α_4_ loop is responsive to chemical intermediates in the desulfurase reaction of EcSufS. A host of results identifies R119 on EcSufE as critical to catalytic turnover, possibly with a role in the EcSufS/EcSufE release step. When compared to SufS homologs, the mobility of the loops appears to be a unique mechanism for catalysis in type II cysteine desulfurase systems.

## Experimental Procedures

### Protein expression and purification

Plasmids containing genes encoding EcSufS (P77444) and EcSufE (P76194) were transformed into BL21(DE3)Δ*suf E. coli* cells, which lack the genomic copy of the *suf* operon. EcSufS was expressed from two different vectors. Expression from a pET44 vector resulted in a *C*-terminal his-tagged EcSufS protein and was used for wildtype and variants with substitutions on the α16 helix. Expression from a pET21 vector resulted in an untagged version of EcSufS and was used for crystallography experiments and in conjunction with the characterization of EcSufE variants. Control activity assays show no difference in activity between the tagged and untagged EcSufS enzymes (Figure S8). All EcSufE enzymes were produced in an untagged version from pET21. QuikChange PCR was used to introduce all mutations discussed in this study using the oligonucleotide primers shown in Table S2. Plasmids harboring mutations were sequenced (Eurofins Genomics, Louisville, KY) to ensure the integrity of the mutated gene.

EcSufS and EcSufE enzymes were expressed via IPTG induction and purified as previously described.(16) His-tagged EcSufS variants from the pET44 vector were expressed via ZYP5052 auto-induction media.(23) A 400 mL culture was prepared with 100 µg/mL ampicillin and incubated at 25 °C for ∼24 hr shaking at 280 rpm. Cells were harvested by centrifugation after reaching saturation (A600 ≥ 10) and stored at −80 °C. To begin purification, cell pellets containing his-tagged EcSufS were resuspended in lysis buffer containing 20 mM MOPS pH 7.5, 300 mM NaCl, 20 mM imidazole, 0.02 mg/mL DNase, 1 mM PMSF, and 5 mM MgCl_2_. Resuspended cells were lysed via sonication and centrifuged at 20,000 ×g for 30 mins at 4 °C to obtain clarified lysate and pellet out cell debris. The clarified lysate containing his-tagged SufS protein was then purified using Ni^2+^ HisTrap HP column with a linear gradient of 20 mM MOPS pH 7.5, 300 mM NaCl, and 20 mM imidazole to 20 mM MOPS pH 7.5, 300 mM NaCl, and 500 mM imidazole. Fractions containing his-tagged SufS protein were analyzed using 12% SDS-PAGE and dialyzed in 20 mM MOPS pH 7.5, 300 mM NaCl to remove the imidazole. Dialyzed his-tagged SufS proteins were then concentrated and stored at −80 °C with 10% glycerol for further use. Concentrations for EcSufS were based on PLP quantification at 388 nm in 0.1 M NaOH (ε_388_ = 6600 M^−1^ cm^−1^).(24) EcSufE concentrations were determined using a calculated ε_280_ extinction coefficient of 20,970 M^−1^ cm^−1^.

### Determining the X-ray structure of the EcSufS/EcSufE complex

To generate a EcSufS/EcSufE complex for crystallization, 230 μM SufS, 230 μM SufE (C51A, E107C) and 2 mM Cys were combined and briefly incubated at room temperature. One μL of protein complex was mixed with 2 μL of crystallization solution containing 24.5-31.5% w/v PEG2000 MME and 0.1 M KSCN in a sitting drop vapor diffusion tray at 20°C. Crystals, which appeared after several days, were cryo-protected by the stepwise addition of mother liquor supplemented with PEG400 to a final concentration of 30% v/v and plunge frozen in liquid nitrogen. Diffraction data were collected at the NE-CAT 24-ID-E beamline of the Advanced Photon Source using a 0.2 sec exposure time per image, 0.979 Å X-rays with 10.2% transmittance and 0.2° oscillation per image. Reflection data were reduced with XDS. Subsequent analysis of the images in Mosflm (as implemented in CCP4i2) indicated anisotropic diffraction. The XDS reflection file was therefore truncated in resolution along the a-axis to 4.2 Å with reflections included to 3.3 Å along b and c using the aniso_cutoff script available at https://wiki.uni-konstanz.de/xds/index.php/Aniso_cutoff.

Structure solution proceeded by determining phases. Molecular replacement was performed using Phaser software and coordinates for the SufS-SufE complex derived from a Rosetta computational model we previously reported but modified to contain one copy of SufS (monomer) and one copy of SufE.(16) The Phaser solution contained two copies of SufS (a homodimer) and two copies of SufE, one bound to each of the SufS active sites. Refinement of the solution was conducted iteratively using Phenix and interactive manual model building in Coot. Rigid body and TLS groups, one group for each polypeptide chain, were used. Molprobity was used to assess the quality of the final coordinates. Statistics regarding structure solution and refinement are presented in Table S1.

### NDA-borate alanine-detection assay

EcSufS can be assayed by following formation of the alanine product using the fluorescence-based naphthalene dicarboxaldehyde (NDA) labeling reaction.(7, 25) In this reaction, NDA reacts with the primary amine of alanine to create a fluorescent adduct that can be detected by HPLC chromatography or multi-well plate reader. Standard assay conditions were 100 mM MOPS pH 8.0, 150 mM NaCl, 2 mM tris(2-carboxyethyl)phosphine (TCEP), 0.25-1 mM EcSufS, 0-5 µM EcSufE, and varying amounts of cysteine (0-500 µM). Robust enzyme activity is followed by a plate-reader assay. For this assay the reaction is initiated by addition of cysteine to the reaction mix at 25 °C. At various time points, 50 µL aliquots are removed and quenched with 5 µL 10% trichloroacetic acid. The samples are labelled by adding 500 µL of an NDA-labelling mix (10 mM borate pH 9.0, 2 mM potassium cyanide, and 0.2 mM NDA) followed by 20 min incubation in the dark. Following the labeling reaction, the fluorescence of each sample was measured using a BioTek Synergy2 multi-well plate reader (Ex 390 nm, Em 440 nm). An alanine standard curve was used to convert fluorescence values into concentration of product. For EcSufS/EcSufE variants that exhibited severe decreases in activity, the more sensitive HPLC-based assay was utilized as previously described.(16)

### Fluorescence Polarization Assay

Fluorescence polarization assays were used to measure the affinity between EcSufS and EcSufE as previously described.(16) Briefly, EcSufE variants were introduced as additional substitutions into a previously constructed C51A/E107C EcSufE variant. This triple variant was labeled by incubation with 500 µM BODIPY-FL maleimide for 4 hr at 25 °C in the dark. After incubation, unreacted dye was removed using a PD-10 desalting column. Labeling efficiency was determined by comparison of the label concentration (ε_505_ = 80,000 M^−1^ cm^−1^) and the EcSufE concentration (ε_280_ = 20,970 M^−1^ cm^−1^) and was between 60-80%. Reaction conditions of 100 nM labeled EcSufE, 50 mM MOPS, pH = 8.0, 150 mM NaCl, and 0.1 g/ml bovine serum albumin were used for titrations with EcSufS (0.1-20 mM). The assay mix was then incubated at 25 °C for 30 min and then transferred to black 96-well plates. Fluorescence polarization (Ex 480 nm, Em 520 nm) was measured using a BioTek Synergy2 multiwell plate reader.

### S75 Size-Exclusion Chromatography

To confirm the quaternary structure and investigate the possibility of dimer formation, purified SufE proteins (wildtype, R119A, and R119K) were subjected to analytical size-exclusion chromatography (SEC) using an S75 10/300 column (Cytiva). The protein samples were diluted to 4 mg/mL in 50 mM MOPS pH 8.0, 150 mM NaCl buffer and 100 μL of each protein variant (wildtype, R119A, and R119K SufE) were loaded onto the S75 column operating at a constant flow rate of 50 mM MOPS pH 8.0, 150 mM NaCl. The chromatograms obtained were analyzed to determine the elution volumes corresponding to the peaks of the SufE protein variants and molecular weight was evaluated based on calibration standards.

### In vivo cluster assembly assay

Biological Fe-S cluster assembly by the SUF pathway can be assessed by overexpressing the operon (*sufABCDSE*) from a pBAD plasmid. Cells with a functional SUF pathway are black/grey colored. Both R119A and R119K SufE substitutions were generated in a pBAD*sufABCDSE* plasmid and confirmed by DNA sequencing. Mutated plasmids were then transformed into Top10 *E. coli* cells. A 100 mL growth was started by 1:100 dilution of overnight growth in LB with 100 μg/mL Amp. Cells were grown until OD600 value of 0.6 and induced with 0.2% arabinose for protein expression. After growth for 3 hours, cells were pelleted and visually inspected for Fe-S cluster formation.

### 35S radiolabel assay

In a 15 µL reaction volume, 30 µM SufS was incubated with 30 µM wildtype SufE, R119A SufE, or R119K SufE and cysteine solution containing 500 µM cold cysteine and 0.1 µM ^35^S cysteine (0.111 µCi/µL). After 5 minutes, the reaction was quenched with 5 µL 250 mM N-ethylmaleimide. The quenched reaction was subjected to gradient SDS-PAGE and visualized by overnight exposure to a phosphoimager screen.

### Data analysis

Steady state kinetic data was fit to the Michaelis-Menten equation (Eq 1) where *v* is the initial velocity, *E*_t_ is the total enzyme concentration, *k*_cat_ is the maximal turnover number, *S* is the substrate concentration, and *K*_M_ is the Michaelis constant. Fluorescence polarization binding data was fit with equation 2 to determine a *K*_D_ value for the interaction. where *A*_o_ is the polarization in the absence of the ligand, *ΔA* is the total change in polarization, and *K*_D_ is the dissociation constant. Each fluorescence polarization assay was fit individually to equation 2 and the parameters were averaged with propagated error.

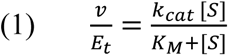

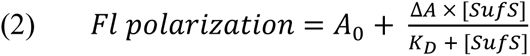

### AlphaFold2-multimer model generation

An AI-generated model of the EcSufS/EcSufE complex was constructed using ColabFold v.1.5.1.(19) The query sequence used the amino acid sequence for two copies of EcSufS and one of EcSufE. Five 3D models were generated, and the highest ranked structure was selected for analysis.

### Sequence-similarity network generation

A representative sequence similarity network(26) was constructed for the IPR003808 domain in the InterPro database (version 92.0) using the Enzyme Function Initiative – Enzyme Similarity Tool (EFI-EST) maintained by the Enzyme Function Initiative (www.enzymefunction.org).(27) The ∼22,000 sequences were reduced to 8,655 representative nodes based on UniRef90 families. The initial network was created at with an alignment cutoff value of 45 or better. The resulting network was downloaded as a Cytoscape readable .xgmml file for visualization.(28)

## Data Availability

All kinetic and biophysical data are contained within the manuscript and supporting information. Structural data have been deposited in the PDB under accession codes 8vbs.

## Supporting Information

This article contains supporting information.

## Author contributions

R.K.G, N.C., M.V., and N.C.G. contributed to the investigation. R.K.G, N.C., M.V., N.C.G., J.A.D., and P.A.F. contributed to the formal analysis. J.A.D. and P.A.F. contributed to the supervision and writing.

## Funding

This work was funded by the National Institutes of Health through grants GM112919 (PAF) and GM142966 (JAD). This work is based upon research conducted at the Northeastern Collaborative Access Team beamlines, which are funded by the National Institute of General Medical Sciences from the National Institutes of Health (P30 GM124165). The Eiger 16M detector on the 24-ID-E beam line is funded by a NIH-ORIP HEI grant (S10OD021527). This research used resources of the Advanced Photon Source, a U.S. Department of Energy (DOE) Office of Science User Facility operated for the DOE Office of Science by Argonne National Laboratory under Contract No. DE-AC02-06CH11357.

## Conflict of Interest

The authors declare that they have no conflicts of interest with the contents of this article.

## Abbreviations

Ala: alanine
Cys: cysteine
DTT: dithiothreitol
EcSufE: SufE from *E. coli*
EcSufS: SufS from *E. coli*
HMM: Hidden-Markov Model
K_D_: dissociation constant
MOPS: 3-morpholinopropane-1-sulfonic acid
NDA: naphthalene 2,3-dicarboxaldehyde
PLP: pyridoxal-5’-phosphate
SSN: sequence similarity network
SUF: sulfur formation pathway
TCEP: tris (2-carboxyethyl)phosphine

## Notes

### Competing Interest Statement

The authors have declared no competing interest.

