## Supplementary material for "The structure of the SufS-SufE complex reveals interactions driving protected persulfide transfer in iron-sulfur cluster biogenesis": SI materials

**Table S1. X-ray data collection and refinement statistics**

| | **$SufS_2$-$SufE_2$ (C51A, E107C)** |
|---|---|
| | **8VBS** |
| **Data collection** | APS 24-ID-E |
| Space group | $P2_12_12$ |
| Cell dimensions | |
| *a, b, c* (Å) | 90.984, 130.134, 100.789 |
| α, β, γ (°) | 90, 90, 90 |
| Wavelength (Å) | 0.979 |
| Resolution (Å)[a] | 79.98 - 3.31 (3.31 - 3.50) |
| $R_{meas}$ (%) | 10.7 (72.9) |
| I/σI | 16.87 (4.28) |
| Completeness (%) | 85.1 (53.6) |
| Redundancy | 11.2 (12.1) |
| $CC_{1/2}$ (%) | 99.9 (96.5) |
| **Refinement** | |
| Resolution (Å) | 60.22 - 3.31 (3.43 - 3.31) |
| No. reflections | 15,754 (934) |
| $R_{work}/R_{free}$ (%) | 23.05 (34.79)/ 27.16 (37.54) |
| No. atoms | 8374 |
| Protein | 8344 |
| Ligand | 30 |
| *B* factors | |
| Protein | 148.12 |
| Ligand | 127.56 |
| R.m.s deviations | |
| Bond lengths (Å) | 0.003 |
| Bond angles (°) | 0.513 |

[a] Values in parentheses are for the highest-resolution shell.

**Table S2. Oligonucleotide mutagenesis primers used in this study.**

| Variant | Primer | Primer Sequence (5'-3') |
|---|---|---|
| H343A SufS | Forward | aatctcggtaaacctatgatgttggc |
| | Reverse | gccaacatcataggtttaccgagatt |
| D346A SufS | Forward | cgagaaaactgccggcgtggtgttta |
| | Reverse | taaacaccacgccggcacttttctcg |
| Y345A SufS | Forward | gagaaaactgccagtgtttaccgaga |
| | Reverse | tctcggtaaacactggcagttttctc |
| N353A SufS | Forward | acgcacagcaatgaaaactgccaaca |
| | Reverse | tgttggcagttttcattgctgtgcgt |
| F350A SufS | Forward | cgtaattatcgagataggcgtggtgt |
| | Reverse | acaccacgcctatctcgataattacg |
| S349A SufS | Forward | gccgtaattatcgatcataggcgtgg |
| | Reverse | ccacgcctatgatcgataattacggc |
| Y354A SufS | Forward | gcacagcaatgcctgccaacatcata |
| | Reverse | tatgatgttggcaggcattgctgtgc |
| T116A SufE | Forward | gtgaacgagatggggcgagatgttgggtgag |
| | Reverse | ctcacccaacatctcgccccatctcgttcac |
| T116S SufE | Forward | acgagatgggctgagatgttgggtgagcgc |
| | Reverse | gcgctcacccaacatctcagcccatctcgt |
| Q121A SufE | Forward | gcttccagacctgctgaacgagatggggtgagatgtt |
| | Reverse | aacatctcaccccatctcgttcagcaggtctggaagc |
| Q121E SufE | Forward | gcttccagaccctctgaacgagatggggtgagatg |
| | Reverse | catctcaccccatctcgttcagagggtctggaagc |
| R119A SufE | Forward | gcttccagaccttgtgaagcagatggggtgagatgttg |
| | Reverse | caacatctcaccccatctgcttcacaaggtctggaagc |
| R119K SufE | Forward | tccagaccttgtgacttagatggggtgagatgttgggtgagc |
| | Reverse | gctcacccaacatctcaccccatctaagtcacaaggtctgga |
| C51A SufE | Forward | gcccagagtcaggtgtggattg |
| | Reverse | gccctgaatgctattttgtgg |
| E107C SufE | Forward | tgcaaaatggcgctcacccaac |
| | Reverse | aaaccacggacgacatcg |

**Figure S1. SDS-PAGE analysis of purified EcSufS and EcSufE variants.** EcSufS variants were run on a 12% SDS-PAGE gel. EcSufE variants were run on a 17% SDS-PAGE gel.

**SufS**

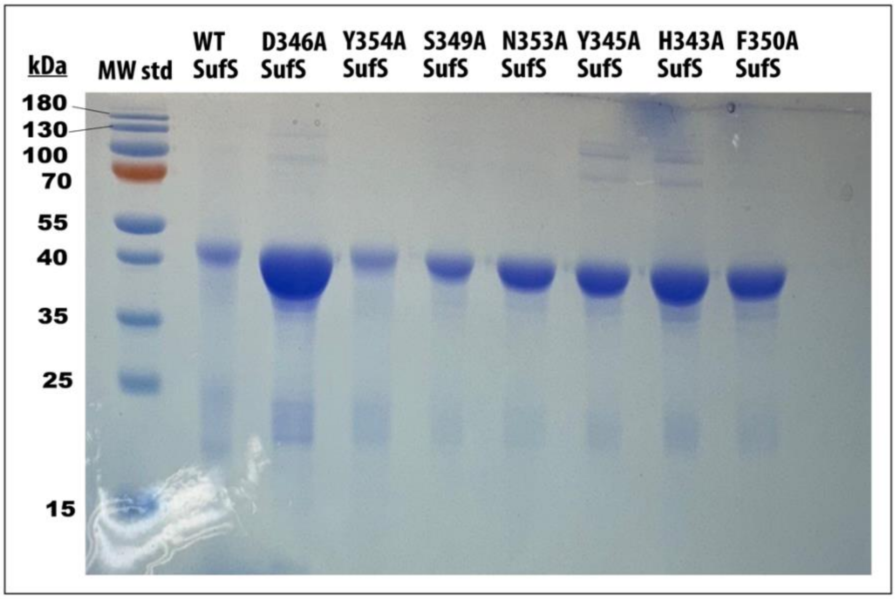

**SufE**

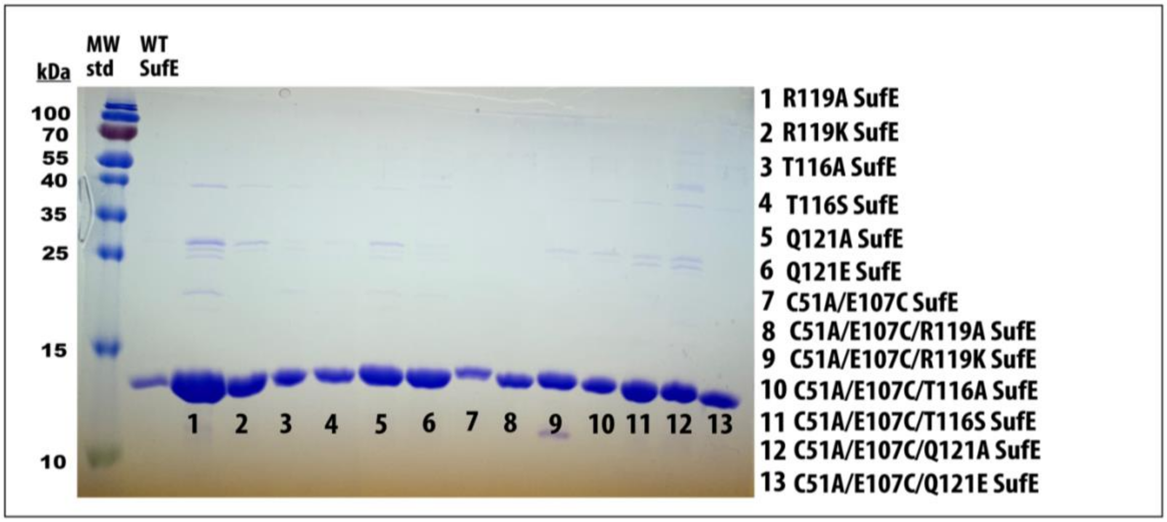

**Figure S2. Circular dichroism spectroscopy of EcSufS and EcSufE variants.** Spectra were acquired in the far-UV region (190-280 nm) at 25°C using a spectropolarimeter (J-1500 CD Spectrometer) for EcSufS (A), EcSufE (B), and C51A/E107C EcSufE (C) variants. Each spectrum is an average of five scans and was baseline corrected using a 50 mM phosphate buffer pH 8.0 as reference. The spectral data was normalized by area to eliminate the changes in each spectrum due to changes in protein concentrations and enable an effective way of comparison of the protein shapes. For the final concentration, protein samples were diluted to 0.2 mg/ml using 50 mM phosphate buffer pH 8.0 to minimize the interference of 150 mM NaCl with the CD spectra in the far-UV region (190-280 nm).

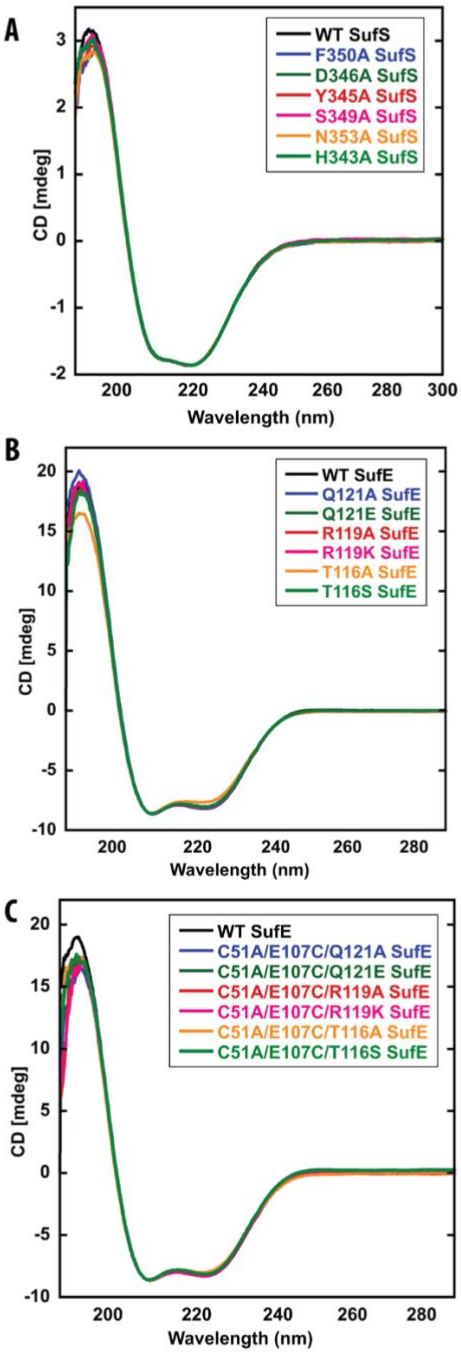

**Figure S3. Fluorescence polarization binding assays for EcSufS variants in the absence of cysteine.** Experimental conditions and analysis described in Materials and Methods section.

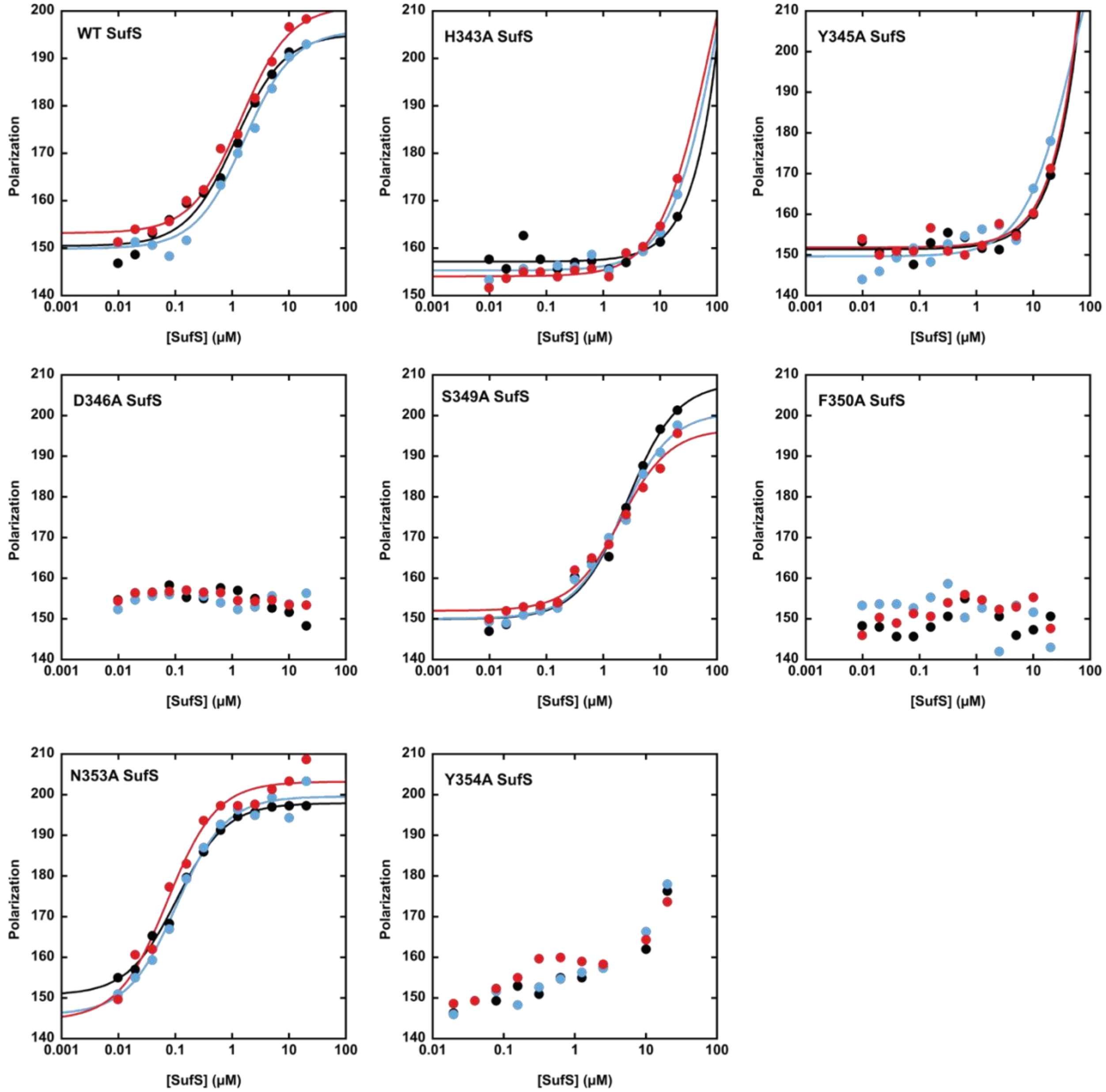

**Figure S4. Fluorescence polarization binding assays for EcSufE variants in the absence of cysteine.** Experimental conditions and analysis described in Materials and Methods section.

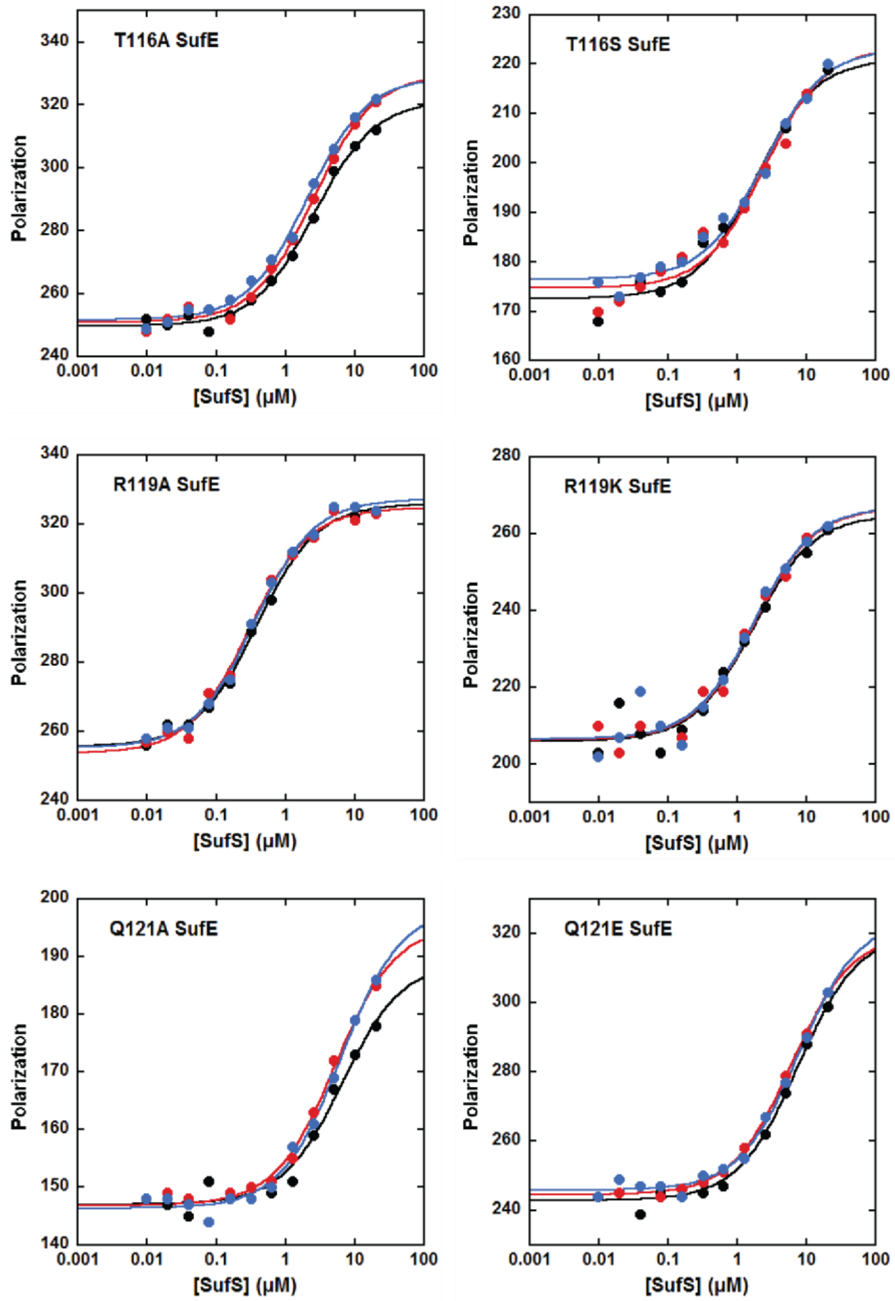

**Figure S5. Fluorescence polarization binding assays for EcSufE variants in the presence of 500 µM cysteine.** Experimental conditions and analysis described in Materials and Methods section.

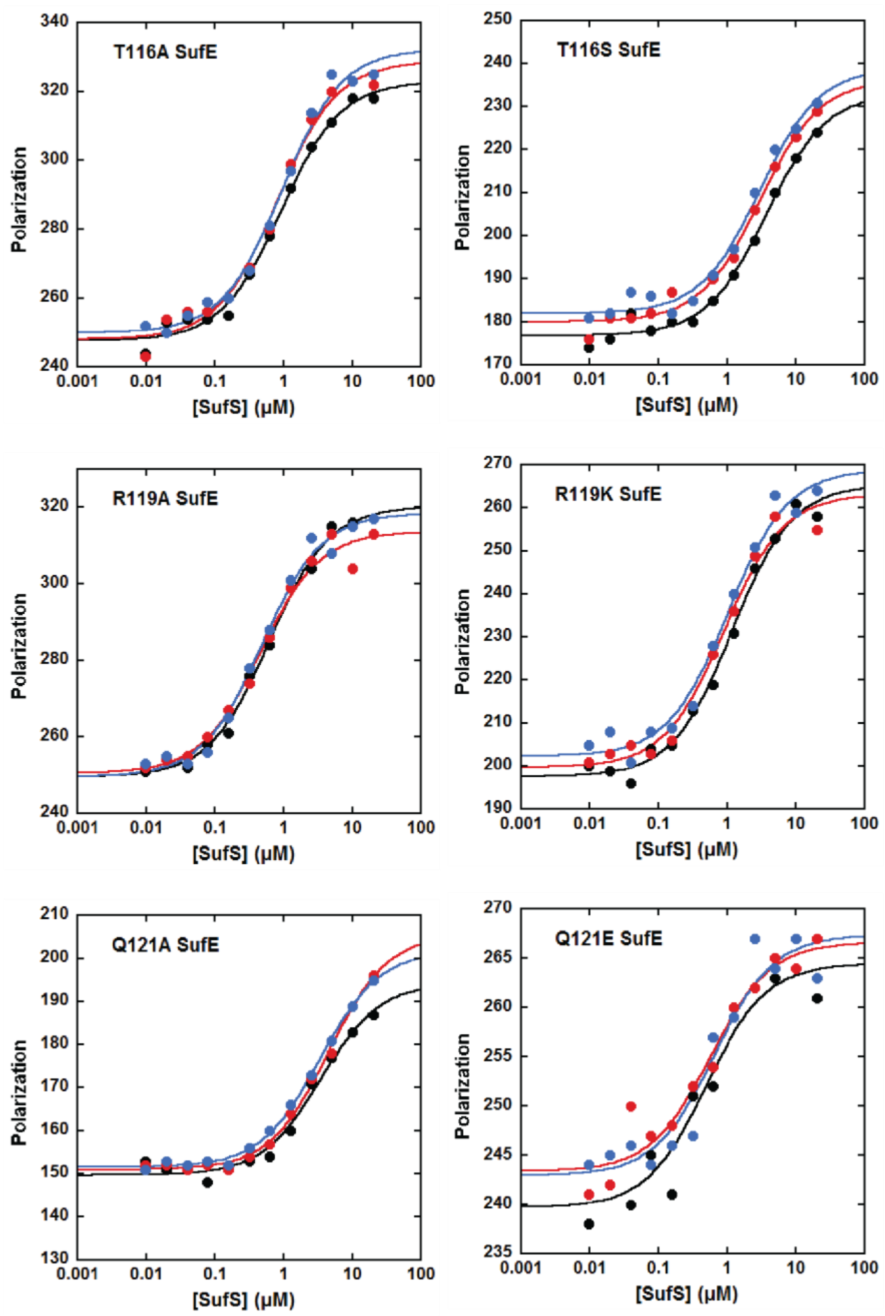

**Figure S6. Structural interactions of R119 in 1mzg.** Dimeric structure is shown with tan and blue monomers. Close of up the active site shows interactions made by R119 in the dimeric crystal structure.

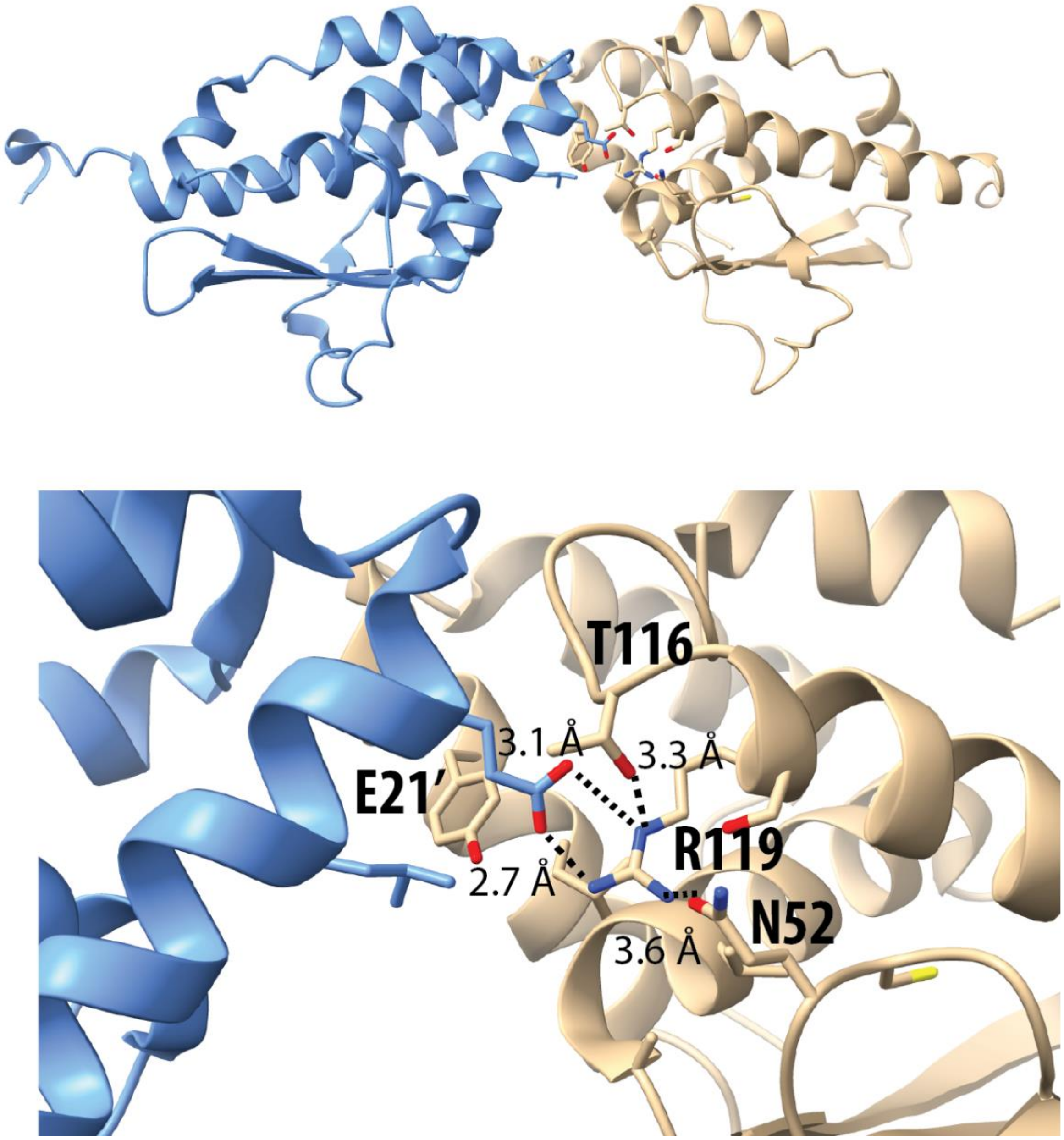

Figure S7. Multiple sequence alignment for the extracted IPR003808 domain from selected SufE-like sequences. 42 sequences from various clusters of the SSN shown in Figure 6. Three sequences were selected from the 14 largest clusters and aligned with MAFFT. Coloring is shown in CLUSTAL scheme for residues at 50% conservation.

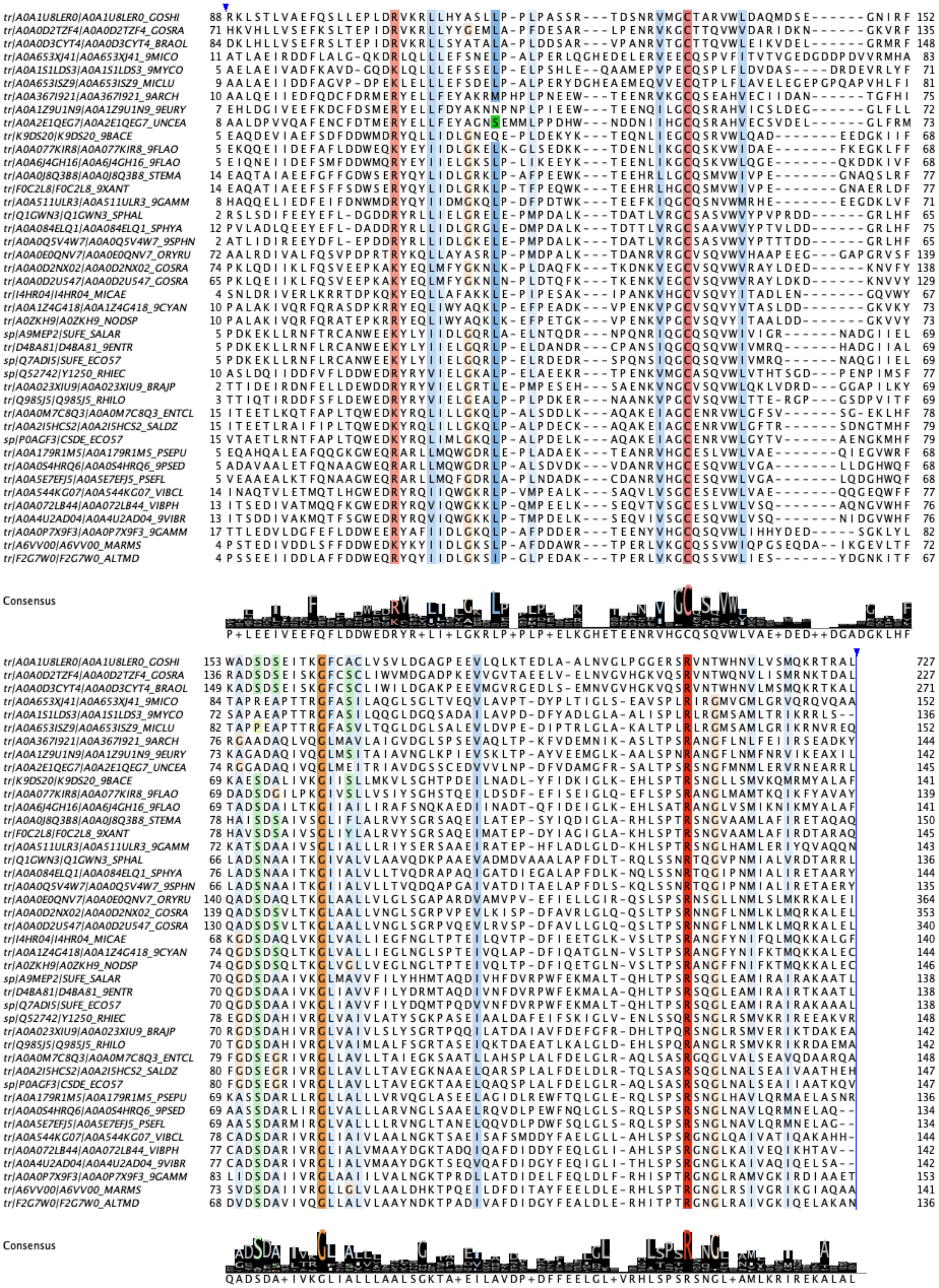

**Figure S8. Michaelis-Menten curve for native and his-tagged EcSufS.** Alanine production was measured by plate reader-based alanine-detection assay. Data are shown for rates of alanine formation for WT SufS (*blue squares*) and WT SufS-HisTag (*black circle*). Reaction conditions are 50 mM MOPS (pH 8.0), 150 mM NaCl, 2 mM TCEP, 500 µM L-cysteine, and 0.2 µM SufS. Initial velocities were determined by varying SufE concentration up to 3 µM. Solid and dotted lines are produced by a fit to Eq 1.

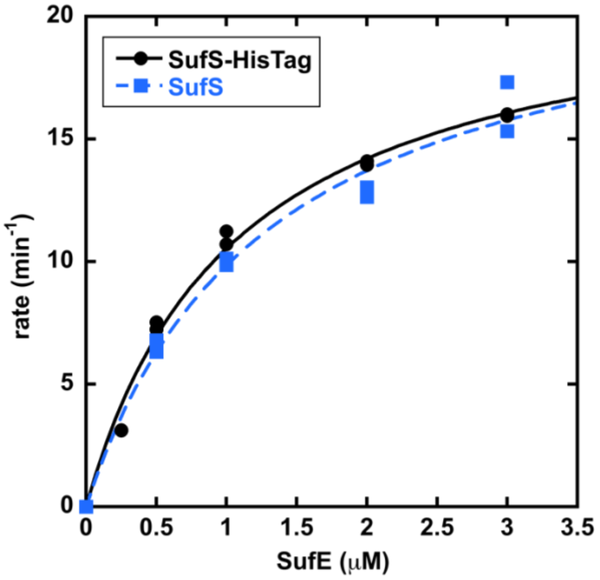
